# Capturing the Hierarchically Assorted Protein-protein Interaction Modules of Mammalian Cell

**DOI:** 10.1101/2024.09.30.615776

**Authors:** Shuaijian Dai, Yage Zhang, Weichuan Yu, Ning Li

## Abstract

Proteins are organized into modules by both functions and physical interactions within compartments of an eukaryotic cell. The *in vivo* chemical crosslinking mass spectrometry (XL-MS) data collected from organelles, the whole cells and tissues are able to provide unique information about both protein-protein interaction (PPI) and the intensity of PPI. In the present study, we have retrieved 55,982 crosslinked peptides (XL-peptides) from the XL-MS databases, out of which 6,356 *in vivo* PPIs were identified. Introduction of the MONET software into analysis of 4,526 hetero PPIs revealed a total of 402 protein modules, including 15, 58 and 163 stable protein complex(s), condensate-forming protein module(s) and intrinsically disordered region (IDR)-containing protein module(s), respectively. The application of ChatGPT in analysis of these modules unexpectedly identified 4 vesicle-related modules. Together, these modules were assorted into 6 communities (module of modules) and 3 systems (module of communities) hierarchically. The bioinformatic analysis found that the three systems are corresponding to three compartments of eukaryotic cell: nuclei, mitochondria, endoplasmic reticulum (ER), respectively. This study presents a novel and comprehensive biomolecular modulome of a mammalian cell, which captures putative protein compositions of protein complexes, protein condensates and vesicles and provides a hierarchical protein organization and function within compartments of mammalian cell.

## Introduction

One of the research goals in life sciences is to correlate the biological phenomena to the behavior of such biomolecules as proteins, nucleic acids, and other small molecules. A good example of this approach is to identify the genetic basis that controls the protein synthesis and interaction underlying the phenotypic variations. Despite the extensive success of this approach, it is still difficult to attribute a complex biological function to an individual molecule. In contrast, most of the biological functions, ranging from DNA replication, transcription, translation to complex signaling cascades, arise from a group or a module of biomolecular components that interact with each other both spatially and temporally in a highly organized and regulated manner. For example, the histone acetyltransferase works in concert with de-acetylation event to regulate the histone acetylation level, which is generally associated with chromatin relaxation, leading to gene activation and expression with the help of transcription factors and RNA polymerase (1, 2). The electron transport chain producing energy in the form of ATP on inner mitochondrial membrane is another example, which consists of several cooperating protein complexes, including NADH dehydrogenases, succinate dehydrogenase, cytochrome c reductase, cytochrome c oxidase and ATP synthase (3, 4). Therefore, the recognition of these structural or functional blocks (also called modules) are critical in understating the biological behavior, which cannot be easily predicted by studying the discrete biochemical and molecular properties of isolated biomolecules. Moreover, these modules are not necessarily being fixed or rigid as they can be quantitatively regulated or switched on and off by cell signals (5). Higher-level functions can be achieved by connecting the interacting modules together to form a hieratically structure of biomolecules (5, 6).

Over the past decade, there has been a significant development in the field of modulomic analysis dedicated to extracting and building modules from complex biomolecular interaction networks. An example is the MoNetFamily web server, which leverages PPI data sourced from the Comprehensive Resource of Mammalian protein complexes database (CORUM 2.0) (7) to explore modules, module-module interaction networks (8), and their variations in module organization (9). ModulOmics is another bioinformatics tool that identifies functionally connected modules that were enriched with cancer driver genes by integrating multiple Omics data types, such as PPI connectivity, DNA mutual exclusivity, transcriptional co-regulation, and co-expression (10). Furthermore, the MTGO (Module detection via Topological information and GO knowledge) tool analyzes PPI networks based on both topological properties and biological knowledge to assemble modules effectively (11). In conjunction with these tools, the multi-layer modularity optimization algorithm (M1 algorithm) embedded in the MONET toolbox (12) has been demonstrated successfully in identifying both the disease-associated modules from proteomic data (13) and the hierarchically organized protein-protein interaction modules within nucleome (6). These studies signify the efficiency of the M1 algorithm in extracting modules of biological functions from PPI networks and in constructing higher order structure and organization of protein modules.

To support the identification and analysis of modules, it is essential to construct large-scale and reliable interaction network of interacting biomolecules (14). In recent years, numbers of PPI data repositories, such as STRING and BioGrid, have been established to record protein connectivity data originating from diverse technologies and research laboratories (15, 16). While these resources have significantly enriched our knowledge on PPIs, these conventional molecular technologies used to map PPIs often face limitations in capturing transient or weak interactions and providing quantitative information on these interactions (6). With the advent of cross-linking mass spectrometry (XL-MS) biotechnology, which has emerged as a useful tool in the field of interactome, it enables the measurement of protein interactions occurred within organelles, cells, or tissues at or near native physiological conditions (17–24). Moreover, XL-MS facilitates the quantitation of PPI abundance by correlating interactions with their corresponding peptide spectrum matches (PSMs), providing valuable insights into the stoichiometry and dynamics of protein modules (6, 17). By integrating the capabilities of XL-MS-based quantitative PPI networks with the M1 algorithm from the MONET toolbox, our previous research uncovered a comprehensive view of the hierarchically assorted liquid nucleome, which included identification of Protein-Protein Interaction Modules (NPIMs) that are associated with protein complexes, nuclear bodies, and protein condensates, along with higher-order communities (modules of modules) that are intricately linked with genomic and nucleolar regions within nuclei (6). Inspired by that successful work, we hereby extended our modulomic analysis pipeline into those *in vivo* XL-MS data derived from organelles, cells, and tissues of mammalian organisms. This Omic approach allowed us to investigate the hierarchical architecture of PPI modules at the whole cell level.

## Experimental procedures

### Collection of XL-MS datasets

The MS raw data of cleavable XL-MS datasets of mammalian cell at *in cell, in tissue* and *in organello* level were obtained from ProteomeXchange Consortium with identifiers of PXD012788, PXD035844, PXD007513, PXD015751, PXD010317, PXD015160, PXD006816 and PXD007673.

### Data analysis and processing

The human proteome database for PXD012788, PXD035844, PXD007513 and PXD015751 was obtained from Uniprot (downloaded on April 7^th^, 2023), containing 20619 sequences. The mouse mitochondrial proteome database for PXD006816 was obtained MitoCarta3.0 (25), containing 1511 sequences. The mouse synapse proteome database for PXD010317 and PXD015160 was downloaded from their original paper (22), containing 5133 sequence entries. The mouse heart proteome database for PXD007673 was downloaded from its original paper (24), containing 4854 target entries.

The MS raw data collected were converted and deconvoluted using Hardklor (26, 27). For *in-silico* protein digestion, the variable modifications were set as oxidation on “M”. The Carbamidomethyl on “C” was set as the fixed modification. The enzyme was set as trypsin. For the crosslinking searching, the tolerance for MS and MS/MS searching were 10 ppm and 20 ppm, respectively. The short-chain monoisotopic mass of the DSSO crosslinker for PXD035844, PXD007513, PXD010317, PXD015160, PXD006816 and PXD014584 was set as 54.0106 Da, while the heavy-chain monoisotopic mass was set as 85.9826 and 103.9932. The short- and heavy-chain monoisotopic mass of the Alkyne-A-DSBSO crosslinker for PXD012788 was set as 54.0106 Da and 236.0177 Da, respectively. The short-chain monoisotopic mass of the BDP-NHP crosslinker for PXD015751 and PXD007673 was set as 197.0324 Da, while the heavy-chain monoisotopic mass was set as 948.4375 Da and 964.4324 Da. The Precursor selection tolerance was 0.05 Da. The fragment type was set as “CID/HCD” for the XL-MS datasets only containing “CID/HCD” MS/MS spectrums and that were set as “CID/HCD” and “ETD” for the datasets containing both “CID/HCD” and “ETD” spectrums. The false discovery rates of the output were calculated by the Percolator (version 3.02) (28, 29) and only the PSMs with q-value ≤ 0.01 was collected.

### Orthologous protein group assignment

Each protein database (human proteome database, mouse mitochondrial proteome database, mouse synapse proteome database and mouse heart proteome database) was searched against EggNOG mammalian orthogroup HMMs using eggnog-mapper (30). The sets of proteins from either human or mouse proteome assigned to the same orthogroup HMM were considered to belong to the same ortholog. The sets of protein-protein interactions assigned to the same ortholog-ortholog interactions were considered as the same protein ortholog-ortholog interactions, and their abundance were calculated as the sum of peptide spectrum matches (PSMs) of the protein-protein interactions (6).

### Protein ortholog interaction normalization

After the PPIs of each dataset were combined into protein ortholog-ortholog interactions, the abundance of PPIs in each dataset were normalized by median normalization using the following equation:

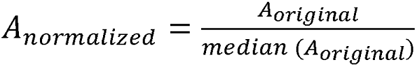

where the A_orginal_ and A_normalized_ stands for the ortholog interaction before and after median normalization. The final abundance of each interaction was calculated as sum of normalized abundance in each dataset.

### Construction of the hierarchically assorted modules of protein-protein interaction

The human and mouse PPIs were first converted into protein ortholog-ortholog interactions. Subsequently, the protein ortholog-ortholog interactions (also called PPI throughout) and their abundance were subsequently processed using R to fit the input format of the MONET toolbox (12, 31). According to the graph theory (32), the two interacting orthologs and the connection between the two orthologs of a ortholog-ortholog interaction were defined as nodes and an edge, respectively. The number of connections on an ortholog node was defined as ortholog degree. The abundance of PPI was calculated as the PSM count(s) of crosslinked peptide(s) that corresponded to an PPI. Both the PPIs (**edges**) and their corresponding abundance were incorporated by the M1 algorithm of the MONET toolbox to construct the modules, communities and systems. The formation of a module required a minimum of two protein orthologs (or called components or **nodes**). Moreover, the entire constituent components of a module could be converted to a single ***converging node***, and the interaction and the corresponding abundance between the two interacting modules can be converted to ***hybrid edge***, respectively. Consequently, module-module interaction (MMI, or called hybrid edge, defines a higher level of edge existing in between two interacting modules). The abundance of MMI (hybrid edge) was the sum of the abundance of PPIs derived from all interacting components between these two modules. Again, both MMI and the abundance of MMI were further incorporated by the M1 algorithm of the MONET toolbox to produce a module of modules, consequently defined as a community. Similarly, the community-community interactions (CCI) were defined as a higher level of interaction existed in between two interacting communities, which is determined by the MMIs among the modules in these two communities. The abundance of CCI was the sum of MMI abundance derived from all interacting modules between these two communities. Both CCI and the abundance of CCI were further incorporated by the M1 algorithm of the MONET toolbox to produce a module of Community (module of module of module), which was consequently defined as a System.

The graph index, system-community-module-component (*x-y-z-w*), indicates the position of a protein or a biomolecule *w* within the System *x*. Those modules that failed to be integrated into a community were defined as the ungrouped (the graph index of this class of module was defined as 0). The diagrams of Module, Community and System of protein orthologs were depicted using Cytoscape (33).

### Protein complex analysis for modules

The well-documented human protein complexes were collected from the Complex Portal (containing 1,715 protein complexes) (34). The protein IDs in each protein complex were transferred into mammalian protein orthologs using the method described in **Orthogroup assignment** section. If the number of components of a protein complex had an overlapping equal to or larger than 50% of that of a module, then the module was defined as a captured protein complex.

### Condensate-forming and IDR-containing protein analysis from modules

The well-reported condensate forming proteins were collected from both the literature and the PhaSePro database (35). The prediction of intrinsic disordered region (IDR) containing proteins was performed MobiDB (36). The proteins having the IDR score in MobiDB ≥ 0.5 was defined as an IDR-containing protein. Next, the protein IDs were converted into mammalian protein orthologs as described in **Orthogroup assignment** section to make them comparable with protein orthologs in each module.

### ChatGPT-assisted function determination for modules

The function and subcellular localization of modules was determined by ChatGPT-3.5 (01 March 2023 version) using R. The R scripts took proteins and their corresponding degree as input, and automatically generated prompt message using the following prompt message with the basic prompt strategy:

’This is a table indicating a protein module. Each row containing the protein name and protein degree. The degree indicates the importance of protein in the module. Use one sentence to describe the main *biological function* of this module. The table is: \n ModuleInfo’

’This is a table indicating a protein module. Each row containing the protein name and protein degree. The degree indicates the importance of protein in the module. Use one word to describe the main *subcellular localization* of this module. The table is: \n ModuleInfo’.

The ModuleInfo here was a table that containing the name and degree information of proteins within a module. Name and degree of a protein was joined by tab (\t), and different proteins are separated by the newline character (\n). Then, the GPT-3.5 was queried using the generated prompt message through OpenAI API (gpt-3.5-turbo-0301), and the returned information was parsed as output.

### Gene Ontology analysis

The Gene Ontology (GO) analysis of the protein orthologs was performed using the R package gprofiler2 (37). The GO annotation database for GO analysis was extracted from the result file of eggnog-mapper (30). The fisher exact test was performed for testing the enrichment of each GO term. The results of the GO analysis were visualized using the R package pheamap and ComplexHeatmap.

### Generation of Random Modules and Communities

The Erdős–Rényi model (38) was applied to generate 100 random protein-protein interaction network with equivalent nodes and edges of the ortholog PPI network in this study. The modulomic analysis (described in section **Construction of the hierarchically assorted modules of protein-protein interaction**) was performed on the random PPI networks to generate modules, communities and systems. The degree of components, number of modules, number of communities and number of systems were analyzed using R.

## Results

### Collection and analysis of XL-MS data in repositories

We applied the modulomic analysis to XL-MS datasets to construct the hierarchically assorted protein-protein interaction modules of mammalian cells. By doing this, seven MS-cleavable XL-MS datasets of mammalian cells (*Homo sapiens* and *Mus musculus*) were downloaded from the ProteomeXchange (39) with the identifier of PXD012788, PXD035844, PXD007513, PXD015751, PXD010317, PXD015160, PXD006816 and PXD007673, followed by XL-peptides searching against target and decoy proteome database using mXlinkX (6, 40, 41), see Experimental procedures for details). As a results, 55,982 repeatable XL-peptides (or called crosslinks) with FDR ≤ 0.01 were collected, which contained 19,439 inter-crosslinks and 36543 intra-crosslinks (Figure 1A; Supplemental Table S1a). The crosslinks were then combined into protein-protein interactions (PPIs) which further converted into 6,356 PPIs (see Experimental procedures for details), containing 4,526 hetero and 1,830 homo PPIs (Figure 1B; Supplemental Table S1b). Majority (3,981 out of 4,526) of the hetero PPIs contained only one type of crosslink. The correlation between the protein interaction and number of crosslinks were found to obey the Zipf’s law with a slope of −2.451 (Figure 1C; Supplemental Table S1c), suggesting an overall distribution of crosslinks identified from XL-MS datastes. The abundance (counts of PSMs) of hetero PPIs of difference datasets (Supplemental Table S1b-c) was consequently normalized using median normalization method and combined into a weighted mammalian ortholog interactions network, containing 3,377 nodes (or proteins) and 4,526 edges (or PPIs, Figure 1D; Supplemental Table S1c). A further investigation revealed that there were 1,133, 416, 144, 712 and 1,243 proteins localized in Nucleus, Mitochondria, Endoplasmic reticulum (ER), Cytosol and other subcellular localizations, respectively, determined using the The Human Protein Atlas annotation (one protein could have muti-localization; Figure 1E; Supplemental Table S1d; Thul et al., 2017). In addition, the degree distribution of 3,377 proteins also followed the Zipf’s law with a slope of −1.798 (Figure 1F; Supplemental Table S1d), indicating an overall distribution of mammalian PPIs identified by XL-MS. Moreover, the Pearson correlation coefficient between the degree and abundance of orthologs was 0.38, showing the highly interactive proteins were not necessarily to be high abundant (Figure 1G).

**Figure 1.**
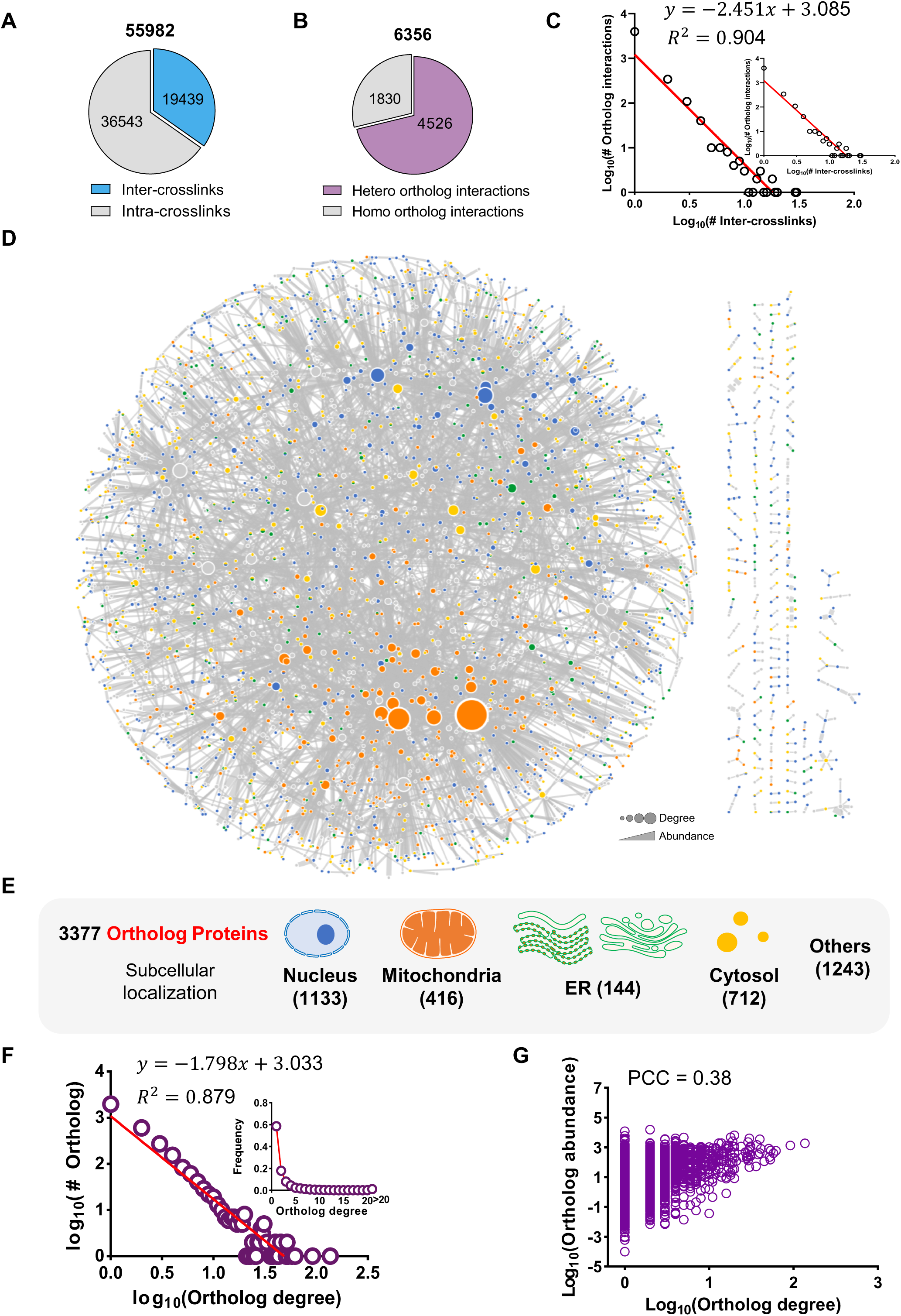
Interactome profile of XL-MS datasets (The figure is related to supplemental figure S1 and Supplemental Table S1). (A) Pie chart represents the distribution of repeatable inter- (blue; 19,439 crosslinks) and intra-crosslinks (grey; 36,543 crosslinks) with FDR ≤ 0.01 identified from the seven XL-MS datasets of mammalian cells (Supplemental Table S1a). (B) Pie chart represents the distribution of hetero- (purple; 4,526 interactions) and homo-ortholog interaction (grey; 1,830 interactions) combined from repeatable crosslinks with FDR ≤ 0.01 (Supplemental Table S1b). (C) The power law equation of the fitting curve (left-bottom panel) describes the correlation in between the total number of hetero ortholog interaction (4,526) and the specific number of crosslinks within an ortholog interaction (19,439 in total). R2 represents the coefficient of the curve determination. The line diagram (right-top panel) shows the distribution of number of crosslinks of ortholog interactions (Supplemental Table S1b-c). (D) Diagram represents the ortholog interaction network. The node and edge indicate the ortholog protein and ortholog interaction, respectively. The size of the node represents the ortholog protein degree. The blue, orange, green, yellow and grey node represents nuclear, mitochondrial, Endoplasmic reticulum (ER), cytosol and other protein, respectively. The thickness of the edge represents the abundance of ortholog interaction (Supplemental Table S1c-d). (E) Diagram shows the subcellular localization of 3,377 ortholog proteins based on the data deposited at The Human Protein Atlas. The subcellular localization of ortholog proteins is classified into five groups which are Nucleus, Mitochondria, Endoplasmic reticulum (ER), Cytosol as well as others (Supplemental Table S1d). (F) The power law equation of the fitting curve (right-top panel) describes the correlation in between the total number of ortholog protein (3,377) and the specific number of degree of an ortholog protein. R2 represents the coefficient of the curve determination. Line diagram (left-bottom panel) shows the distribution of degree of 3,377 ortholog proteins derived 4526 hetero-ortholog interaction (Supplemental Table S1d). (G) Diagram represents the correlation between degree and abundance of ortholog proteins. The abundance of proteins are obtained from Supplemental Table S1d. The PCC stands for the Pearson Correlation Coefficient.

### Construction of hierarchically architecture of mammalian cells

To build modules from the PPI data, we utilized the M1 algorithm of the MONET toolbox (12) by considering both the interactions and their corresponding abundance (first round of modulomic analysis; Supplemental Table S1c), resulting in the identification of 402 individual modules (≥ 2 components per module; Figure 2A; Supplemental Table S2a). The higher-order structure of the modules was revealed by a second round of modulomic analysis. To do this, the nodes edges and the abundance of protein interaction within a module were merged as converging nodes, hybrid edges and abundance of module-module interaction (MMI), respectively. The MMI data was thereafter analyzed by M1 algorithm, resulted in module of modules, *i.e.* communities. A total of 241 modules were constructed into six distinct communities, Community 1 (C1), Community 2 (C2), Community 3 (C3), Community 4 (C4), Community 5 (C5) and Community 6 (C6), comprising of 58, 54, 40, 37, 32 and 20 modules, respectively. The resting 161 modules remained ungrouped. Similar to the community construction, the community-community interaction (CCI) network was also constructed and analyzed to generate module of community, *i.e.* Systems by the third round of modulomic analysis, to depict the architecture and organization of communities. Consequently, three systems, System 1 (S1), System 2 (S2) and System 3 (S3), were identified and they were made of C2 and C3, C1 and C5 as well as C4 and C6, respectively (Figure 2A; Supplemental Table S2a). Through the modulomic analysis, we were able to construct a hierarchically assorted modulome of mammalian cell, which is comprised of systems, communities, modules and protein node (Figure 2A; Supplemental Table S2a). The graph index, system-community-module-component (x-y-z-w), was used to indicate the position of a component (protein node) w.

**Figure 2.**
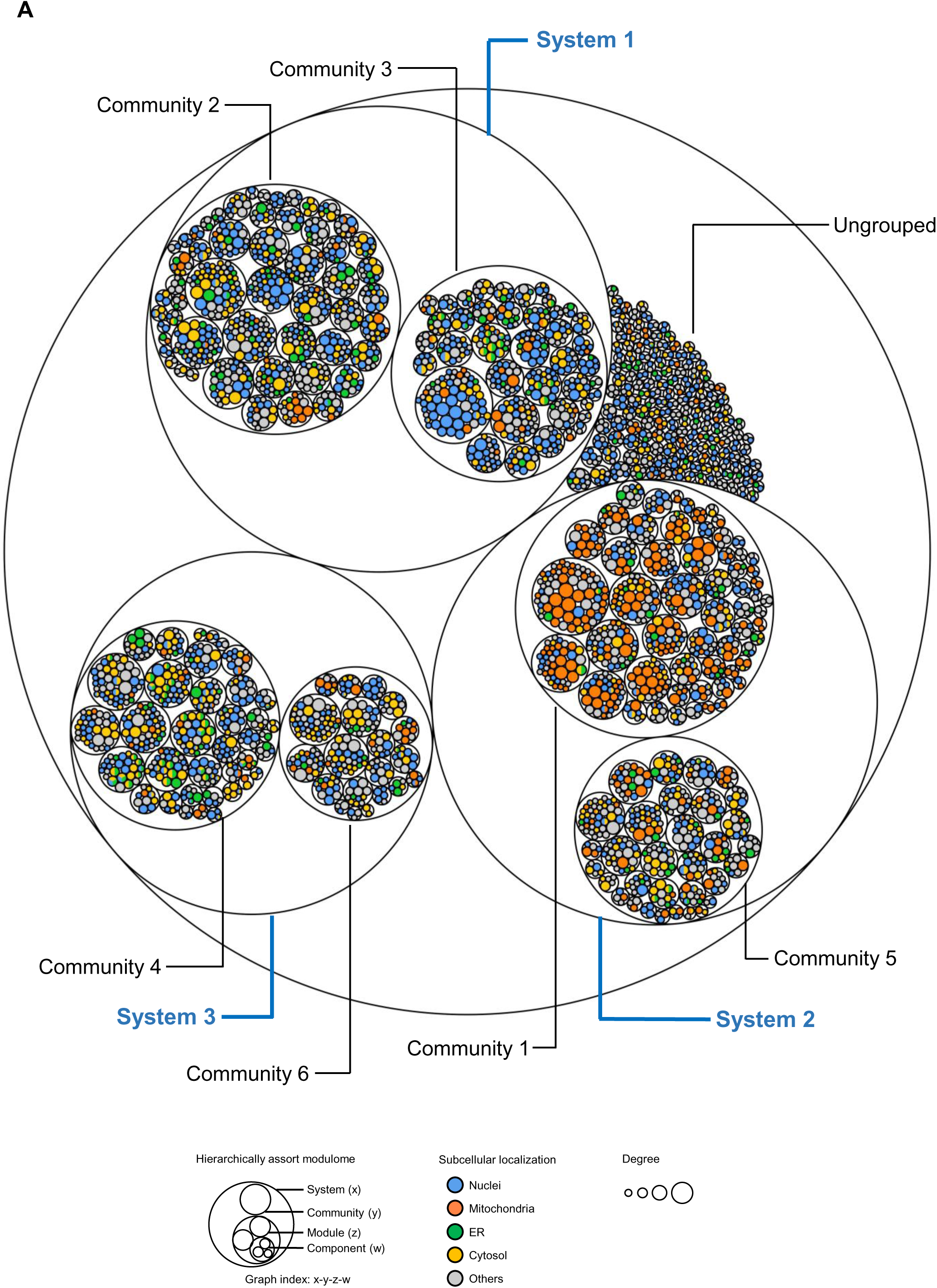
Hierarchically assorted modulome of mammalian cells (The figure is related to supplemental Table S2). (A) Circle plot denotes the hierarchically assorted modulome containing system, community, modules and protein components, generated by multiple rounds of modulomic analysis of ortholog interaction network. Node denotes the ortholog proteins. The size of node denotes the protein degree. The blue, orange, green, yellow and grey node represents nuclear, mitochondrial, ER, cytosol and other ortholog protein, respectively. The outline of grouped nodes (ortholog protein components) denotes a module. The outline of grouped modules denotes a community. The outline of grouped community denotes a system. Ungrouped are those modules that fail to be integrated into a community. The data is listed in Supplemental Table S2a.

### Identification and characterization of the functions of protein modules

In total, 402 modules were constructed during the first round of modulomic analysis. Of these, 167 modules had a minimum size of two, while the largest module comprised 67 components (Figure 3A; Supplemental Table S2a). The distribution of the size of modules also followed the Zipf’ law (Figure 3A; Supplemental Table S2a). As we proposed previously, these modules were likely to represent protein complexes and condensates (6). Therefore, we compared the modules with the well-document protein complexes deposited at Complex Portal (34) and condensates that were deposited at both the literature and the PhaSePro database (35). We observed that 15 and 58 modules overlapped with the well-reported protein complexes and condensates, respectively, further supporting our original findings (Figure 3B; Supplemental Table S2a). Furthermore, we predicted the intrinsic disordered region (IDR) of ortholog proteins using the MobiDB program, resulting in 163 modules containing one component possessing the predicted IDR domain (Figure 3B; Supplemental Table S2a). We compared the properties of known condensate forming proteins with other proteins in the modules and found that they were likely to have higher level of degree and were more likely to be classified into modules (Figure 3C-D; Supplemental Table S2a).

**Figure 3.**
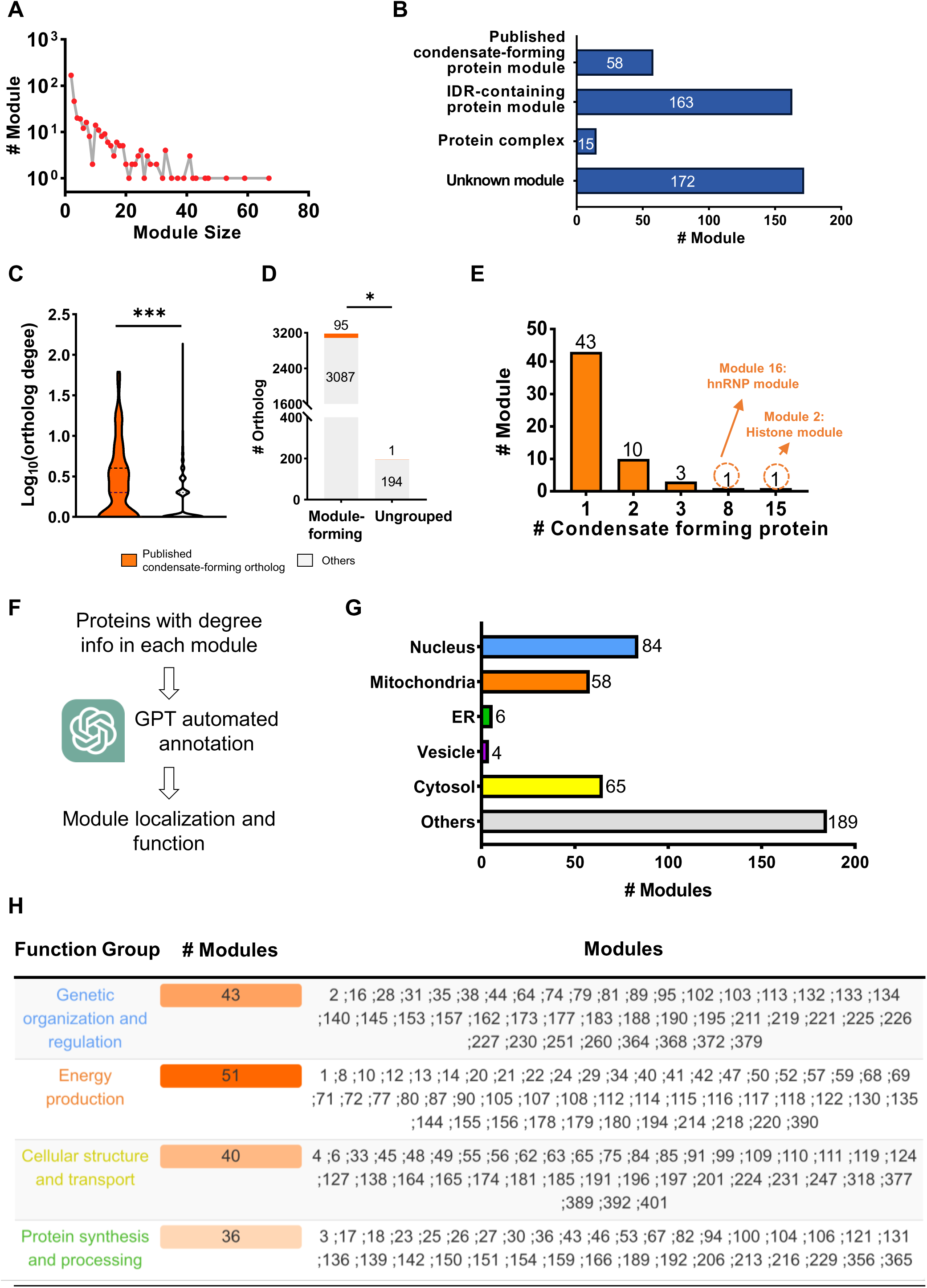
Bioinformatic characterization of modules (The figure is related to supplemental Table S2). (A) Line plot (left-bottom panel) denotes the distribution of module size (Supplemental Table S2a). (B) Bar chart represents the characterization of modules. The modules are classified into four groups, published condensate-forming protein module (58), predicted IDR-containing protein module (163), protein complex module (15) and uncharacterized module (172) (Supplemental Table S2a). (C) Violin plot shows the degree of published condensate-forming proteins (orange) and that of other ortholog proteins (grey). Fisher exact test was applied. The n.s, *, **, and *** stands for p > 0.05, p < 0.05, p < 0.01 and p < 0.001, respectively. The data is listed in Supplemental Table S2a. (D) Bar chart denotes the distribution of number of literature-annotated condensate-forming ortholog protein within (module-forming) and outside (ungrouped) modules. Ungrouped are those ortholog proteins that fail to be integrated into a module. The number in each bar denotes the number of ortholog protein in each category. The orange and light grey color stands for literature-annotated condensate-forming and other ortholog protein, respectively. Fisher exact test was applied. The n.s, *, **, and *** stands for p > 0.05, p < 0.05, p < 0.01 and p < 0.001, respectively. The data is listed in Supplemental Table S2a. (E) Bar chart denotes the distribution of the number of published condensate-forming protein in modules. The modules are classified into five groups that contain 1 (43), 2 (10), 3 (3), 8 (1) and 15 (1) published condensate-forming proteins (Supplemental Table S2a). (F) Pipeline of ChatGPT-assisted function annotation of modules. (G) Bar chart denotes the distribution of subcellular localization of modules annotated by ChatGPT (Supplemental Table S2b). (H) Table containing the grouping information of ChatGPT-assisted function annotation. The functions of modules were classified into four groups which were genetic organization and regulation, energy production, protein synthesis and processing as well as cellular structure and transporting. The column 1, 2, 3 stands for the class of function, number of modules within the function group and the exact number of modules, respectively (Supplemental Table S2b).

A subsequent analysis of condensate modules revealed 43 modules contained one condensate-forming protein, while module 1-3-2 and module 1-2-16 were composed of 15 and 8 distinct condensate-forming proteins, respectively (Figure 3E; Supplemental Table S2a). Module 1-3-2 was found to be a putative histone module, containing fifteen well-documented condensate proteins of nucleosome histone components, including histone H1/H5, histone H2A, histone H2B, and histone H3. This suggests that it may be correlated with chromatin formation, which has been reported to have an intrinsic liquid-liquid phase separation (LLPS) capacity (Supplemental Table S2a). Similarly, module 1-2-16 was identified as a possible heterogeneous nuclear ribonucleoproteins (hnRNPs) module, containing six well-documented condensate proteins of hnRNP complex (43). These proteins are involved in mRNA splicing, transcription, and translation regulation, and their structure contains intrinsically disordered regions (IDRs), which can form condensates through the interaction between IDRs (Supplemental Table S2a). Module 8 contained the condensate-forming protein FAST kinase domains 3 (FASTD3), which is associated with mitochondrial RNA granules (MRG) to form fluid condensates (44). Module 12-1-46 contained mitochondrial import receptor subunit TOM70, which is involved in the localized protein condensation of cytosolic proteins on the mitochondria surface (45). Module 3-4-30 was composed of Zonula Occluden 1 (ZO-1) protein, which was reported to form cytoplasmic droplets through multivalent interaction to drive the formation of tight junctions (46, 47). Moreover, two condensate-forming proteins, microtubule-associated protein Tau and cytoplasmic protein NCK1, were colocalized in module 3-6-92, suggesting its association with cell motility.

To enhance our understanding of the subcellular localization and functional roles of the 402 identified protein modules, we utilized ChatGPT, an advanced AI language model, to predict these attributes based on integrated datasets (Figure 3F). The localization predictions indicated that 84 modules are in the nucleus, 58 in the mitochondria, 6 in the ER, 4 in vesicles, 65 in the cytosol, and 189 in other subcellular locations (Figure 3G; Supplemental Table S2b). Notably, we identified four well-documented vesicle-associated modules, specifically module 45, 73, 185, and 231 (Supplemental Table S2b). Module 1-2-73 includes SEC22B and STX12, key components of the SNARE complex involved in cargo transport between the ER and Golgi apparatus, as well as in the endosomal-lysosomal pathway. Similarly, module 1-2-185 contains COPB1, ARCN1, and COPA, essential for the coatomer protein complex I (COPI), which is critical in vesicle formation and trafficking within the Golgi apparatus and retrograde transport to the ER.

Regarding biological functions, 172 modules could not be assigned due to insufficient component information. The remaining modules were categorized into four functional groups (Figure 3H). The genetic organization and regulation group includes 43 modules involved in gene expression regulation, DNA replication, and other nuclear processes. The energy production group, comprising 51 modules, is engaged in metabolic pathways such as ATP production, the citric acid cycle, and oxidative phosphorylation. The cellular structure and transportation group, with 40 modules, likely maintains cellular integrity and facilitates intracellular transport, including vesicle trafficking, organelle movement, and cytoskeletal organization. Lastly, 36 modules are associated with protein synthesis and processing, involved in protein production, folding, modification, and degradation, ensuring proper protein function and turnover.

### Characterization of functions of communities and systems

As for the communities and systems, it was interesting to find that modules containing the well-documented condensate-forming proteins were significantly enriched in the group of modules that form communities as comparing with that in the ungrouped modules (Figure 4A-B; Supplemental Table S2a). Consequently, we did the similar investigation to each community and found that C2 and C3 had the top2 largest proportion of published condensate-forming proteins containing modules (Figure 4C; Supplemental Table S2a). Moreover, to characterize the biological role of each community and system, the gene ontology (GO) analysis was performed on the highly interactive components of the communities of each system (6). For the C2 and C3 derived from S1, the analysis revealed that proteins were largely located at chromatin. Chromosome, nucleosome and nuclear lumen by the cellular component, involved in chromatin assembly or disassembly, chromatin silencing, chromatin organization and nucleosome organization by biological process and eventually specialized in nucleic acid binding, chromatin DNA binding and nucleosomal DNA binding by molecular function (Figure 4D-E, Supplemental Figure S3A; Supplemental Table S3a-c). Therefore, the S1 was defined as a Nuclei System. Similarly, the cellular component results showed that the S2, containing C1 and C3, were highly enriched in mitochondrion-related terms such as mitochondrion, mitochondrial envelope, mitochondrial membrane protein complex, mitochondrial matrix, mitochondrial respirasome and respiratory chain complex (Figure 4D; Supplemental Table S3a-c), suggesting it was associated with Mitochondrial System, which could be further confirmed with the biological process enrichment in generation of precursor metabolites and energy, ATP synthesis coupled electron transport, oxidative phosphorylation, electron transport chain and ATP metabolic process (Figure 4E; Supplemental Table S3a-c) as well as molecular function enrichment in oxidoreductase activity, NADH dehydrogenase activity and electron transfer activity (Supplemental Figure S2A; Supplemental Table S3a-c). Moreover, the S3 was defined as ER System as the highly interactive proteins were enriched in ribosome and actin cytoskeleton cellular component, peptide biosynthetic process, translation, translational initiation, nuclear-transcribed mRNA catabolic process, protein transport and protein targeting to ER for biological process as well as structural constituent of ribosome and RNA binding of molecular function (Figure 4D-E, Supplemental Figure S2A; Supplemental Table S3a-c).

**Figure 4.**
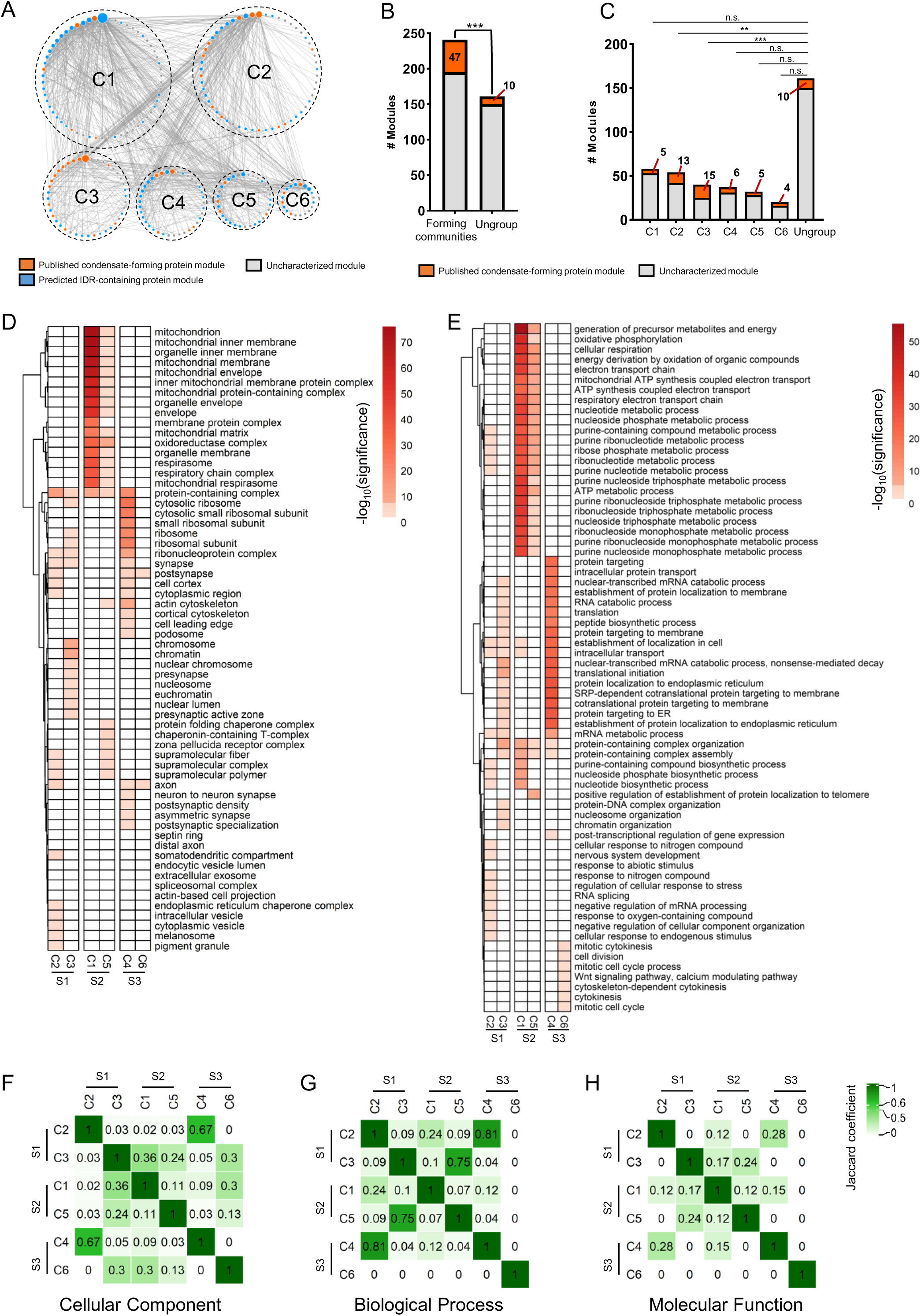
Characterization of Communities and Systems Constructed from Ortholog Protein Interaction Network. (The figure is related to supplemental figure S2 and supplemental Table S2-3). (A) Network diagram represents seven communities present in mammalian cell. The node and edge represents the module and the interaction among modules, which is defined as module-module interaction (MMI), respectively. The higher order of modules (module of module) is defined as community. The size of the node represents the degree of module. The orange, light blue and light grey node represents the published condensate-forming, the predicted IDR-containing and uncharacterized module, respectively. The thickness of line represents the abundance of MMI. C1 – C6 stands for community 1 - 6. The data is listed in Supplemental Table S2a. (B) Bar chart denotes the distribution of number of literature-annotated condensate-forming protein module within (community-forming) and outside (ungrouped) communities. Ungrouped are those modules that fail to be integrated into a community. The orange and grey color stands for modules containing and not containing published condensate-forming proteins, respectively. The number in each bar denotes the number of modules containing the condensate-forming protein. Fisher exact test was applied. The n.s, *, **, and *** stands for p > 0.05, p < 0.05, p < 0.01 and p < 0.001, respectively. The data is listed in Supplemental Table S2a. (C) Bar chart denotes the distribution of number of literature-annotated condensate-forming protein module in each community as well as that of modules outside communities. The orange and grey color stands for modules containing and not containing published condensate-forming proteins, respectively. The number in each bar denotes the number of modules containing the condensate-forming protein. C1 – C6 stands for community 1 - 6. Ungrouped are those modules that fail to be integrated into a community. Fisher exact test was applied. The n.s, *, **, and *** stands for p > 0.05, p < 0.05, p < 0.01 and p < 0.001, respectively. The data is listed in Supplemental Table S2a. (D - E) The heatmap of cellular component (D) and biological process (E) enrichment analysis of the ortholog proteins obtained each community. The top 10% most interactive proteins (highest ortholog degree) in each community were used for the analysis. The communities are organized by systems. Row represents the name of GO term while column marks the different system and community of ortholog proteins. The color represents the negative number of base 10 logarithm of each GO term. S1 - S3 stands for system 1 – 3. C1 – C6 stands for community 1 - 6. The data is listed in Supplemental Table S3a-c. (F - H) The heatmap shows the comparison of cellular component (F), biological process (G) and molecular function (H) enrichment analysis of ortholog proteins derived from seven communities which are organized into systems. The top 10% most interactive proteins (highest ortholog degree) in each community were used for the analysis. The similarity among the communities was evaluated by Jaccard Coefficient (JC) of GO terms. The green color of the heatmap stands for the level of Jaccard Coefficient. S1 - S3 stands for system 1 – 3. C1 – C6 stands for community 1 - 6. The data is listed in Supplemental Table S3a-c.

Furthermore, our analysis of the subcellular localization of modules by ChatGPT within systems S1, S2, and S3 revealed similar patterns with GO analys**is** (Figure 4D-E, Supplemental Figure S2A-B; Supplemental Table S3a-c). Nucleus-localized modules were predominantly found in S1, mitochondria-localized modules were enriched in S2, and all ER-related modules were present in S3 (Supplemental Figure S2B). These AI-produced findings reinforce our characterization of the three systems as representing distinct cellular compartments: nuclei, mitochondria and ER. In addition, the subcellular localization of highly interactive proteins annotated by the Human Protein Atlas in each system also aligns with the GO analysis (Supplementary Figure S2C). In system S1, there was a significant enrichment of nuclear proteins, while mitochondrial and endoplasmic reticulum (ER) proteins were notably reduced. Conversely, in system S2, mitochondrial proteins were significantly up-regulated, whereas nuclear and ER proteins were markedly down-regulated. For system S3, ER proteins were significantly enriched, and nuclear and mitochondrial proteins were suppressed.

To study the correlation among the three systems, the comparison of the enrichment of GO terms in each community organized by systems were investigated, revealing that the overall association between two systems were relatively low (Figure 4F-H). It was interesting to find that the correlation of biological processes between C3 in S1 and C4 in S3 was high which could be the result of the communication between transcription and translation activities (Figure 4H).

Taken together, the integration of XL-MS interactome and MONET-based modulomic analysis had allowed us to organize the individual protein components (nodes) into a hierarchically assorted modulome, containing structure of systems communities, modules and components. The hierarchical structure could hereby be represented by graph index (see Experimental procedures for details). In addition, the well-separated system of proteins, S1, S2 and S3 could be topologically and biologically confined to Nuclei, Mitochondria and ER, respectively (Figure 5; Supplemental Table S2a).

**Figure 5.**
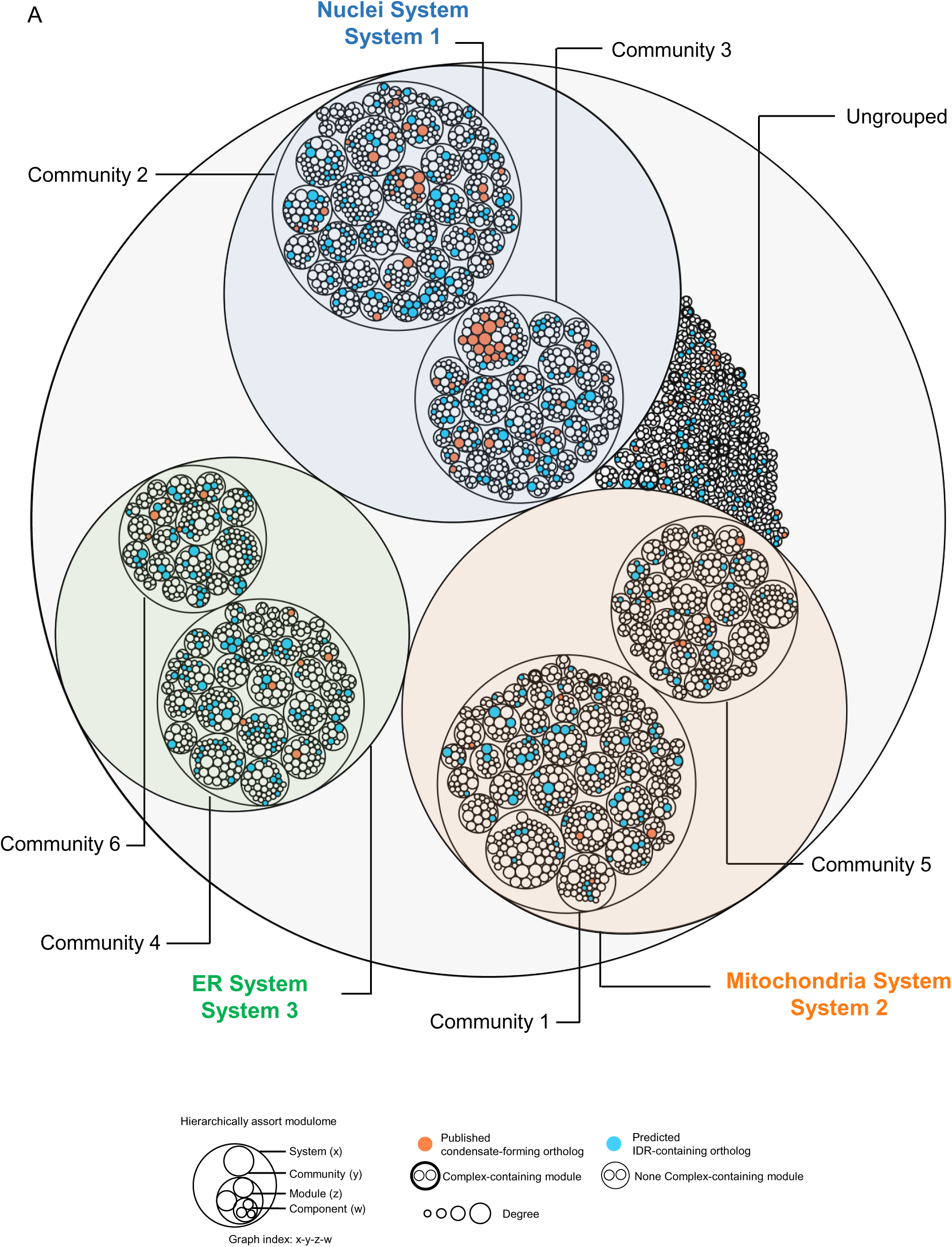
Overview of hierarchically assorted modulome of mammalian cells. (The figure is related to supplemental Table S2). (A) Circle plot denotes the hierarchically assorted modulome containing system, community, modules and protein components, generated by multiple rounds of modulomic analysis of ortholog interaction network. Circle denotes the ortholog proteins. The size of circle denotes the protein degree. The orange color, blue color, and white color filled circle represents published condensate-forming, IDR-containing and uncharacterized ortholog protein, respectively. The bold outline of circle indicates a module which is overlapped with well-reported protein complex. The outline of grouped circles (ortholog protein components) denotes a module. The outline of grouped modules denotes a community. The outline of grouped community denotes a system. Ungrouped are those modules that fail to be integrated into a community. The biological character of each system is determined by GO enrichment analysis. The data is listed in Supplemental Table S2a.

## Discussion

### Integration of multiple XL-MS datasets of *Homo sapiens* and *Mus musculus*

To construct modulome of mammalian cells, we downloaded available MS-cleavable XL-MS datasets of *Homo sapiens* and *Mus musculus* deposited at the ProteinXchange (48). These datasets were further filtered by the biological sample used to generate these data. Only those datasets made by crosslinking of organelles, living cells and tissues were selected to ensure the reliability of the captured *in vivo* or *in organello* PPIs. The mXlinkX was used in this study for the searching of crosslinked peptides as the reliability of this software have been confirmed in our previous study (6). Since the XL-MS datasets were generated by different samples, organisms as well as laboratories, we invented a method to combine these PPIs and normalize their corresponding abundance. In order to combine the PPIs of *Homo sapiens* and *Mus musculus*, we converted the PPIs into mammalian protein orthologs by using eggnog-mapper (30), which had been successfully applied in a pan-plant proteomic study to integrate the proteins from different plant species (49). As for the normalization of abundance of PPIs, we used the median normalization method which was widely used for normalizing gene expression data across microarrays (50) and has been successfully applied in peptide data in LC-MS analysis (51).

### Proper controls used in the modulomic analysis

To ensure our findings were not due to randomness, we first generated 100 random PPI Erdös-Rényi networks, each with the same number of nodes (3,377) and edges (4,526) (Supplemental Figure S3A). The number of modules in these networks varied from 56 to 108, while the number of communities ranged from 2 to 5 (Supplemental Figure S3B-C). Additionally, each random network comprised only one system (Supplemental Figure S3D). These results clearly indicated that the modulome constructed in this study is unlikely to be a consequence of randomness.

In our previous work (6), we proposed that the abundance of PPIs is crucial for modulome construction. To explore this, we created two special networks: an unweighted PPI network with identical protein connections but with all abundances set to 1, and a weighted PPI network with the same connections but random abundance assignments (Supplemental Figures S4, S5; Supplemental Table S4a-b). The unweighted PPI network resulted in 236 modules, 3 communities, and 1 system (Supplemental Figure S4A; Supplemental Table S4c). However, the unweighted PPI network showed very little similarity in module and community structures compared to our study (Supplemental Figure S4B-C; Supplemental Table S4c). Similarly, the randomly weighted PPI network produced 272 modules, 2 communities, and 1 system (Supplemental Figure S5B-C; Supplemental Table S4d), with significant differences in the components of each module and community (Supplemental Figure S5A-C; Supplemental Table S4d). These findings collectively demonstrate the critical role of the *in vivo* PPI abundance in modulomic analysis.

We also considered whether the source of PPIs could influence modulome construction. To test this, we collected PPIs for the 3,377 proteins in this study from STRING database (Supplemental Table S4e). This resulted in a PPI network with 2,316 nodes and 39,196 edges, which we then analyzed by the modulomics workflow (Supplemental Figure S6A-B; Supplemental Table S4e). The resulting modulome comprised 344 modules, 14 communities, and 2 systems (Supplemental Figure S6A-B; Supplemental Table S4f). Comparison with our study’s modulome showed low functional similarity at both the module and community levels (Supplemental Figure S7; Supplemental Table S4f), highlighting the importance of XL-MS-generated PPIs in protein modulome construction.

Furthermore, we examined whether different search engines could affect the results of modulome construction. We summarized an ortholog protein interaction network using the XL-MS results deposited at the original papers (18, 20–24, 52), each employing different search engines and data analysis pipelines. This network contained 3,969 ortholog proteins and 2,476 hetero ortholog interactions (Supplemental Figure S8A-B; Supplemental Table S4g). The modulomic analysis revealed a hierarchically assorted structure with 411 modules, 7 communities, and 3 systems (Supplemental Figure S8C-D; Supplemental Table S4g). Further investigation into the biological roles of these communities and systems identified two systems: the Nuclei system and the Mitochondria system, while the third system’s role remained unclear based on GO analysis (Supplemental Figure S8D; Supplemental Table S4h). These results suggest that although search engines and data analysis pipelines may introduce some variations, we could still obtain the similar higher-order structure of eukaryotic cells, which could be searching engine and data analysis methods independent.

### Biological roles of module, community and system

Previously, we have demonstrated the potential correlation between the modules and protein complexes or protein condensates. In this study, we also found 15 and 58 modules which contains protein complexes and published condensate-forming proteins, respectively (Figure 3B; Supplemental Table S2a), which further supports our previous hypothesis. In fact, the identified known protein complexes could be a confirmation of our module construction. Interestingly, 15 modules contained larger or equal to two condensate-forming proteins (Figure 3E; Supplemental Table S2a), suggesting the module’s potential in depicting the condensate protein composition. For instance, module 1-3-2 was likely to be concentrated with well-studied intrinsic chromatin condensates (53) while module 1-2-16 was possibly made of hnRNP condensates (Figure 3E; Supplemental Table S2a). In addition, two m6A-binding proteins, YTH N6-methyladenosine RNA binding protein 2 (YTHDF2) and YTH N6-methyladenosine RNA binding protein 3 (YTHDF3) were co-identified in module 63. These two proteins have been reported to phase-separate together to form protein droplets (54). As the composition of biomolecular condensates are difficult to be analyzed using traditional biochemistry methods due to the difficulties in isolation of condensates (55). The modulomic analysis pipeline has enabled the discovery of both putative scaffold and client components of a condensate.

The hierarchically structure of modules, communities and systems, were built through modulomic analysis whereas bioinformatic analysis of the highly interactive proteins in each system and community has found that these systems may be potentially correlated with specific organelles or subcellular regions of cells, whereas communities may represent subregions within those organelles or subcellular regions. As for the S1, the C3 of S1 was enriched with histone proteins as well as GO terms in chromatin and DNA binding, suggesting it could be the Genome hub of nucleus. Although the C2 region of S2 could not be definitively classified as a sub-region of the nucleus due to the lack of clear indications in the function enrichment analysis, the co-location of two key nucleolus components, Nucleophosmin 1 (NPM1) and Fibrillarin (FBL), within C2 provides evidence that it may be the nucleolus region of the nucleus. The insignificance of nucleolus-related terms in function enrichment analysis could be the results of lacking enough nucleolus related XL-MS results as the histone proteins in module 1-3-2 has total degree of 252 while the NPM1 in module 1-2-35 only had a degree of 19 and FBL in module 1-2-88 only had a degree of 1 (Supplemental Table S2a). The C1 of S2 was putatively associated with inner membrane and respirasome of mitochondrial according to the significant enrichment of correlated GO terms (Figure 4D-E, Supplemental Figure S2A; Supplemental Table S3a-c). The C5 of S2 was not clearly separated from C1 as their GO enrichment patterns were similar. However, the similar GO terms in C5 were not as significant as that in C1 and C5 had some unique enrichment in cytoskeleton and protein localization (Figure 3D-E). In addition, the proteins with top2 largest degree in C5 were myosin and prohibitin-2 (PHB2), which were function in mitochondrial trafficking (56, 57) and protein import (58), respectively. Myosin is localized in the outer membrane of the mitochondria (57), whereas the C-terminus of PHB2 is exposed to the intermembrane space (59). Thus, C5 could represent the outer membrane and intermembrane space of the mitochondrial and may be responsible for organelle movement and protein importation. Moreover, the C4 in S3 were likely to represent the ribosome region according to the GO analysis (Figure 3D-E, Supplemental Figure S2A) and function in protein translation while the role of C6 was not clear as only few function terms were enriched. Similar to the S1, the biological role of C6 might be determined with the increasing number of XL-MS datasets.

### ChatGPT-assisted characterization of modulome

Characterizing modules using GO and KEGG analysis is challenging due to the lack of diversity in data sources, the need for up-to-date data, the ability to integrate information, and the capability to reduce the interference of noise within the modules. On the contrary, utilizing ChatGPT’s predictive capabilities enabled us to assign subcellular localizations and functional annotations to the 402 modules identified in this study. Notably, ChatGPT accurately pinpointed vesicle-associated modules, highlighting essential components such as SEC22B, STX12, COPB1, ARCN1, and COPA, which play critical roles in vesicle formation and trafficking. Moreover, the functional categorization of these modules by ChatGPT provided valuable insights into four major biological groups: genetic organization and regulation, energy production, cellular structure and transportation, and protein synthesis and processing (Figure 3H; Supplemental Table S2b). In addition, the distribution of module localization across the three systems was consistent with the results of GO analysis, which further support our findings and also illuminate the effectiveness of AI in uncovering the complex biological characteristics within protein modules.

However, it is notable that current AI systems heavily rely on the quality of training and input data, which could lead to inaccuracies in prediction results. Furthermore, interpreting the results is challenging due to the ‘black box’ nature of AI processes. Despite these challenges, the evolution of AI has significantly enhanced our understanding of modular biomolecules, making substantial contributions to the fields of bioinformatics and molecular biology.

### Limitation and potential of modulomic analysis in 4D biomolecular modulome

The power of modulomic analysis pipeline through the combination of XL-MS-based interactome and MONET-based analysis has been demonstrated on both the nucleome level in our previous study (6) and cell level in this study. The hieratically structure of proteins were deciphered and organized in component-module-community-system, with each level possessing its unique biology characteristics. However, the current analysis was limited by the number of crosslinks of *in organello* or *in vivo* XL-MS studies. As more XL-MS studies are conducted, it will become feasible to construct the entire cell graph, as we proposed in our earlier work (6). Additionally, the present modulome represents an average across different cell types at various developmental stages. By incorporating cell-type-specific XL-MS studies, we have the potential to explore the 4D biomolecular modulome at the tissue, organ, and even organism levels (Supplemental Figure S9). This advancement could significantly enhance our overall understanding of biological systems and how biomolecules interact and function, probably transport, in cellular compartments.

## Author Contributions

SJD contributes to data retrieving and modulomic analysis. SJ.D., N.L., YG.Z., and WC.Y. wrote the manuscript; N.L. and YG.Z. contributed to the revision of manuscript and the financial support to this work; N.L. was in charge of project planning, designing, project execution, and communication with collaborators and responsible for distribution of materials and methods integral to the findings.

## Data availability

The XL-MS data used in this manuscript were downloaded from PrteomeXchange with identifiers of PXD012788, PXD035844, PXD007513, PXD015751, PXD010317, PXD015160, PXD006816 and PXD007673.

## Supplemental Data

This article contains supplemental data.

## Declarati1on of interests

No conflict of interest is declared.

## Abbreviations

XL-MS: chemical crosslinking mass spectrometry
PSMs: peptide spectrum matches
NPIMs: Nuclear Protein-Protein Interaction Modules
PPI: protein-protein interaction
MMI: module-module interaction
CCI: community-community interaction
ER: endoplasmic reticulum
LLPS: lipuid-liquid phase separation
IDRs: intrinsically disordered regions
hnRNPs: heterogeneous nuclear ribonucleoproteins
FASTD3: FAST kinase domains 3
MRG: mitochondrial RNA granules
COPI: coatomer protein complex I
YTHDF2: YTH N6-methyladenosine RNA binding protein 2
YTHDF3: YTH N6-methyladenosine RNA binding protein 3
NPM1: Nucleophosmin 1
FBL: Fibrillarin
PHB2: Prohibitin-2

## Reference

1. Bannister, A. J., and Kouzarides, T. (2011) Regulation of chromatin by histone modifications. Cell Research 21, 381–395

2. Peixoto, L., and Abel, T. (2013) The role of histone acetylation in memory formation and cognitive impairments. Neuropsychopharmacology 38, 62–76

3. Pfanner, N., Warscheid, B., and Wiedemann, N. (2019) Mitochondrial proteins: from biogenesis to functional networks. Nature Reviews Molecular Cell Biology 20, 267–284

4. Giacomello, M., Pyakurel, A., Glytsou, C., and Scorrano, L. (2020) The cell biology of mitochondrial membrane dynamics. Nature Reviews Molecular Cell Biology 21, 204–224

5. Hartwell, L. H., Hopfield, J. J., Leibler, S., and Murray, A. W. (1999) From molecular to modular cell biology. Nature 402, 47–52

6. Dai, S., Liu, S., Zhou, C., Yu, F., Zhu, G., Zhang, W., Deng, H., Burlingame, A., Yu, W., Wang, T., and Li, N. (2023) Capturing the hierarchically assorted modules of protein–protein interactions in the organized nucleome. Molecular Plant, 930–961

7. Giurgiu, M., Reinhard, J., Brauner, B., Dunger-Kaltenbach, I., Fobo, G., Frishman, G., Montrone, C., and Ruepp, A. (2019) CORUM: The comprehensive resource of mammalian protein complexes - 2019. Nucleic Acids Research 47, D559–D563

8. Lin, C.-Y., Lin, Y.-W., Yu, S.-W., Lo, Y.-S., and Yang, J.-M. (2012) MoNetFamily: a web server to infer homologous modules and module-module interaction networks in vertebrates. Nucleic acids research 40, W263–70

9. Lin, C. Y., Lee, T. L., Chiu, Y. Y., Lin, Y. W., Lo, Y. S., Lin, C. T., and Yang, J. M. (2015) Module organization and variance in protein-protein interaction networks. Scientific Reports 5, 1–12

10. Silverbush, D., Cristea, S., Yanovich-Arad, G., Geiger, T., Beerenwinkel, N., and Sharan, R. (2019) Simultaneous Integration of Multi-omics Data Improves the Identification of Cancer Driver Modules. Cell Systems 8, 456–466.e5

11. Vella, D., Marini, S., Vitali, F., Di Silvestre, D., Mauri, G., and Bellazzi, R. (2018) MTGO: PPI Network Analysis Via Topological and Functional Module Identification. Scientific Reports 8, 1–13

12. Tomasoni, M., Gómez, S., Crawford, J., Zhang, W., Choobdar, S., Marbach, D., and Bergmann, S. (2020) MONET: A toolbox integrating top-performing methods for network modularization. Bioinformatics 36, 3920–3921

13. Johnson, E. C. B., Carter, E. K., Dammer, E. B., Duong, D. M., Gerasimov, E. S., Liu, Y., Liu, J., Betarbet, R., Ping, L., Yin, L., Serrano, G. E., Beach, T. G., Peng, J., De Jager, P. L., Haroutunian, V., Zhang, B., Gaiteri, C., Bennett, D. A., Gearing, M., Wingo, T. S., Wingo, A. P., Lah, J. J., Levey, A. I., and Seyfried, N. T. (2022) Large-scale deep multi-layer analysis of Alzheimer’s disease brain reveals strong proteomic disease-related changes not observed at the RNA level. Nature Neuroscience 25, 213–225

14. Rodina, A., Xu, C., Digwal, C. S., Joshi, S., Patel, Y., Santhaseela, A. R., Bay, S., Merugu, S., Alam, A., Yan, P., Yang, C., Roychowdhury, T., Panchal, P., Shrestha, L., Kang, Y., Sharma, S., Almodovar, J., Corben, A., Alpaugh, M. L., Modi, S., Guzman, M. L., Fei, T., Taldone, T., Ginsberg, S. D., Erdjument-Bromage, H., Neubert, T. A., Manova-Todorova, K., Tsou, M. F. B., Young, J. C., Wang, T., and Chiosis, G. (2023) Systems-level analyses of protein-protein interaction network dysfunctions via epichaperomics identify cancer-specific mechanisms of stress adaptation. Nature Communications 14,

15. Szklarczyk, D., Gable, A. L., Lyon, D., Junge, A., Wyder, S., Huerta-Cepas, J., Simonovic, M., Doncheva, N. T., Morris, J. H., and Bork, P. (2019) STRING v11: protein–protein association networks with increased coverage, supporting functional discovery in genome-wide experimental datasets. Nucleic acids research 47, D607–D613

16. Oughtred, R., Rust, J., Chang, C., Breitkreutz, B.-J., Stark, C., Willems, A., Boucher, L., Leung, G., Kolas, N., Zhang, F., Dolma, S., Coulombe-Huntington, J., Chatr-aryamontri, A., Dolinski, K., and Tyers, M. (2021) The BioGRID database: A comprehensive biomedical resource of curated protein, genetic, and chemical interactions. Protein Science 30, 187–200

17. Liu, S., Yu, F., Hu, Q., Wang, T., Yu, L., Du, S., Yu, W., and Li, N. (2018) Development of in Planta Chemical Cross-Linking-Based Quantitative Interactomics in Arabidopsis. Journal of Proteome Research 17, 3195–3213

18. Yu, C., Wang, X., Huszagh, A. S., Viner, R., Novitsky, E., Rychnovsky, S. D., and Huang, L. (2019) Probing H(2)O(2)-mediated Structural Dynamics of the Human 26S Proteasome Using Quantitative Cross-linking Mass Spectrometry (QXL-MS). Molecular & cellular proteomicslJ: MCP 18, 954–967

19. Wheat, A., Yu, C., Wang, X., Burke, A. M., Chemmama, I. E., Kaake, R. M., Baker, P., Rychnovsky, S. D., Yang, J., and Huang, L. (2021) Protein interaction landscapes revealed by advanced in vivo cross-linking-mass spectrometry. Proceedings of the National Academy of Sciences of the United States of America 118,

20. Bartolec, T. K., Vázquez-Campos, X., Norman, A., Luong, C., Johnson, M., Payne, R. J., Wilkins, M. R., Mackay, J. P., and Low, J. K. K. (2023) Cross-linking mass spectrometry discovers, evaluates, and corroborates structures and protein–protein interactions in the human cell. Proceedings of the National Academy of Sciences 120, e2219418120

21. Fasci, D., Ingen, H. Van, Scheltema, R. A., and Heck, A. J. R. (2018) Histone interaction landscapes visualized by crosslinking mass spectrometry in intact cell nuclei. Molecular and Cellular Proteomics 17, 2018–2033

22. Gonzalez-Lozano, M. A., Koopmans, F., Sullivan, P. F., Protze, J., Krause, G., Verhage, M., Li, K. W., Liu, F., and Smit, A. B. (2020) Stitching the synapse: Cross-linking mass spectrometry into resolving synaptic protein interactions. Science Advances 6, 1–15

23. Liu, F., Lössl, P., Rabbitts, B. M., Balaban, R. S., and Heck, A. J. R. (2018) The interactome of intact mitochondria by cross-linking mass spectrometry provides evidence for coexisting respiratory supercomplexes. Molecular and Cellular Proteomics 17, 216–232

24. Chavez, J. D., Lee, C. F., Caudal, A., Keller, A., Tian, R., and Bruce, J. E. (2018) Chemical Crosslinking Mass Spectrometry Analysis of Protein Conformations and Supercomplexes in Heart Tissue. Cell Systems 6, 136–141.e5

25. Rath, S., Sharma, R., Gupta, R., Ast, T., Chan, C., Durham, T. J., Goodman, R. P., Grabarek, Z., Haas, M. E., Hung, W. H. W., Joshi, P. R., Jourdain, A. A., Kim, S. H., Kotrys, A. V, Lam, S. S., McCoy, J. G., Meisel, J. D., Miranda, M., Panda, A., Patgiri, A., Rogers, R., Sadre, S., Shah, H., Skinner, O. S., To, T.-L., Walker, M. A., Wang, H., Ward, P. S., Wengrod, J., Yuan, C.-C., Calvo, S. E., and Mootha, V. K. (2021) MitoCarta3.0: an updated mitochondrial proteome now with sub-organelle localization and pathway annotations. Nucleic acids research 49, D1541–D1547

26. Hoopmann, M. R., Finney, G. L., and MacCoss, M. J. (2007) High-speed data reduction, feature detection, and MS/MS spectrum quality assessment of shotgun proteomics data sets using high-resolution mass spectrometry. Analytical chemistry 79, 5620–5632

27. Hoopmann, M. R., MacCoss, M. J., and Moritz, R. L. (2012) Identification of peptide features in precursor spectra using Hardklör and Krönik. Current protocols in bioinformatics Chapter 13, Unit13.18-Unit13.18

28. The, M., MacCoss, M. J., Noble, W. S., and Käll, L. (2016) Fast and Accurate Protein False Discovery Rates on Large-Scale Proteomics Data Sets with Percolator 3.0. Journal of the American Society for Mass Spectrometry 27, 1719–1727

29. Käll, L., Canterbury, J. D., Weston, J., Noble, W. S., and MacCoss, M. J. (2007) Semi-supervised learning for peptide identification from shotgun proteomics datasets. Nature methods 4, 923–925

30. Huerta-Cepas, J., Szklarczyk, D., Heller, D., Hernández-Plaza, A., Forslund, S. K., Cook, H., Mende, D. R., Letunic, I., Rattei, T., Jensen, L. J., von Mering, C., and Bork, P. (2019) eggNOG 5.0: a hierarchical, functionally and phylogenetically annotated orthology resource based on 5090 organisms and 2502 viruses. Nucleic Acids Research 47, D309–D314

31. Arenas, A., Fernández, A., and Gómez, S. (2008) Analysis of the structure of complex networks at different resolution levels. New Journal of Physics 10,

32. Al-Taie, M. Z., and Kadry, S. (2017) Graph theory

33. Smoot, M. E., Ono, K., Ruscheinski, J., Wang, P.-L., and Ideker, T. (2011) Cytoscape 2.8: new features for data integration and network visualization. BIOINFORMATICS APPLICATIONS NOTE 27, 431–43210

34. Meldal, B. H. M., Bye-A-Jee, H., Gajdoš, L., Hammerová, Z., Horáčková, A., Melicher, F., Perfetto, L., Pokorný, D., Lopez, M. R., Türková, A., Wong, E. D., Xie, Z., Casanova, E. B., Del-Toro, N., Koch, M., Porras, P., Hermjakob, H., and Orchard, S. (2019) Complex Portal 2018: Extended content and enhanced visualization tools for macromolecular complexes. Nucleic Acids Research 47, D550–D558

35. Mészáros, B., Erdős, G., Szabó, B., Schád, É., Tantos, Á., Abukhairan, R., Horváth, T., Murvai, N., Kovács, O. P., Kovács, M., Tosatto, S. C. E., Tompa, P., Dosztányi, Z., and Pancsa, R. (2020) PhaSePro: the database of proteins driving liquid–liquid phase separation. Nucleic Acids Research 48, D360–D367

36. Piovesan, D., Necci, M., Escobedo, N., Monzon, A. M., Hatos, A., Mičetić, I., Quaglia, F., Paladin, L., Ramasamy, P., Dosztányi, Z., Vranken, W. F., Davey, N. E., Parisi, G., Fuxreiter, M., and Tosatto, S. C. E. (2021) MobiDB: Intrinsically disordered proteins in 2021. Nucleic Acids Research 49, D361–D367

37. Raudvere, U., Kolberg, L., Kuzmin, I., Arak, T., Adler, P., Peterson, H., and Vilo, J. (2019) G:Profiler: A web server for functional enrichment analysis and conversions of gene lists (2019 update). Nucleic Acids Research 47, W191–W198

38. Erdös, P., and Rényi, A. (1959) On Random Graphs I. Publicationes Mathematicae Debrecen 6, 290

39. Deutsch, E. W., Csordas, A., Sun, Z., Jarnuczak, A., Perez-Riverol, Y., Ternent, T., Campbell, D. S., Bernal-Llinares, M., Okuda, S., Kawano, S., Moritz, R. L., Carver, J. J., Wang, M., Ishihama, Y., Bandeira, N., Hermjakob, H., and Vizcaíno, J. A. (2017) The ProteomeXchange consortium in 2017: supporting the cultural change in proteomics public data deposition. Nucleic Acids Research 45, D1100–D1106

40. Liu, F., Rijkers, D. T. S., Post, H., and Heck, A. J. R. (2015) Proteome-wide profiling of protein assemblies by cross-linking mass spectrometry. Nature Methods 12, 1179–1184

41. Liu, F., Lössl, P., Scheltema, R., Viner, R., and Heck, A. J. R. (2017) Optimized fragmentation schemes and data analysis strategies for proteome-wide cross-link identification. Nature Communications 8,

42. Thul, P. J., Akesson, L., Wiking, M., Mahdessian, D., Geladaki, A., Ait Blal, H., Alm, T., Asplund, A., Björk, L., Breckels, L. M., Bäckström, A., Danielsson, F., Fagerberg, L., Fall, J., Gatto, L., Gnann, C., Hober, S., Hjelmare, M., Johansson, F., Lee, S., Lindskog, C., Mulder, J., Mulvey, C. M., Nilsson, P., Oksvold, P., Rockberg, J., Schutten, R., Schwenk, J. M., Sivertsson, A., Sjöstedt, E., Skogs, M., Stadler, C., Sullivan, D. P., Tegel, H., Winsnes, C., Zhang, C., Zwahlen, M., Mardinoglu, A., Pontén, F., Von Feilitzen, K., Lilley, K. S., Uhlén, M., and Lundberg, E. (2017) A subcellular map of the human proteome. Science 356,

43. Domanski, M., Dedic, E., Pérez, M. E., Cléry, A., Campagne, S., Uldry, A.-C., Braga, S., Heller, M., Rabl, J., Afanasyev, P., Boehringer, D., Nováček, J., Allain, F. T., and Mühlemann, O. (2022) 40S hnRNP particles are a novel class of nuclear biomolecular condensates. Nucleic Acids Research 50, 6300–6312

44. Rey, T., Zaganelli, S., Cuillery, E., Vartholomaiou, E., Croisier, M., Martinou, J.-C., and Manley, S. (2020) Mitochondrial RNA granules are fluid condensates positioned by membrane dynamics. Nature cell biology 22, 1180–1186

45. Liu, Q., Fong, B., Yoo, S., Unruh, J. R., Guo, F., Yu, Z., Chen, J., Si, K., Li, R., and Zhou, C. (2023) Nascent mitochondrial proteins initiate the localized condensation of cytosolic protein aggregates on the mitochondrial surface. Proceedings of the National Academy of Sciences 120, e2300475120

46. Kinoshita, N., Yamamoto, T. S., Yasue, N., Takagi, C., Fujimori, T., and Ueno, N. (2022) Force-dependent remodeling of cytoplasmic ZO-1 condensates contributes to cell-cell adhesion through enhancing tight junctions. iScience 25, 103846

47. Beutel, O., Maraspini, R., Pombo-García, K., Martin-Lemaitre, C., and Honigmann, A. (2019) Phase Separation of Zonula Occludens Proteins Drives Formation of Tight Junctions. Cell 179, 923–936.e11

48. Deutsch, E. W., Bandeira, N., Perez-Riverol, Y., Sharma, V., Carver, J. J., Mendoza, L., Kundu, D. J., Wang, S., Bandla, C., Kamatchinathan, S., Hewapathirana, S., Pullman, B. S., Wertz, J., Sun, Z., Kawano, S., Okuda, S., Watanabe, Y., Maclean, B., Maccoss, M. J., Zhu, Y., Ishihama, Y., and Vizcaíno, J. A. (2023) The ProteomeXchange consortium at 10 years: 2023 update. Nucleic Acids Research 51, D1539–D1548

49. McWhite, C. D., Papoulas, O., Drew, K., Cox, R. M., June, V., Dong, O. X., Kwon, T., Wan, C., Salmi, M. L., Roux, S. J., Browning, K. S., Chen, Z. J., Ronald, P. C., and Marcotte, E. M. (2020) A Pan-plant Protein Complex Map Reveals Deep Conservation and Novel Assemblies. Cell 181, 460–474.e14

50. Yang, Y. H., Dudoit, S., Luu, P., Lin, D. M., Peng, V., Ngai, J., and Speed, T. P. (2002) Normalization for cDNA microarray data: a robust composite method addressing single and multiple slide systematic variation. Nucleic acids research 30, e15

51. Ejigu, B. A., Valkenborg, D., Baggerman, G., Vanaerschot, M., Witters, E., Dujardin, J.-C., Burzykowski, T., and Berg, M. (2013) Evaluation of normalization methods to pave the way towards large-scale LC-MS-based metabolomics profiling experiments. OmicslJ: a journal of integrative biology 17, 473–485

52. Yao, R.-W., Xu, G., Wang, Y., Shan, L., Luan, P.-F., Wang, Y., Wu, M., Yang, L.-Z., Xing, Y.-H., Yang, L., and Chen, L.-L. (2019) Nascent Pre-rRNA Sorting via Phase Separation Drives the Assembly of Dense Fibrillar Components in the Human Nucleolus. Molecular cell 76, 767–783.e11

53. Gibson, B. A., Blaukopf, C., Lou, T., Chen, L., Doolittle, L. K., Finkelstein, I., Narlikar, G. J., Gerlich, D. W., and Rosen, M. K. (2023) In diverse conditions, intrinsic chromatin condensates have liquid-like material properties. Proceedings of the National Academy of Sciences 120, e2218085120

54. Ries, R. J., Zaccara, S., Klein, P., Olarerin-George, A., Namkoong, S., Pickering, B. F., Patil, D. P., Kwak, H., Lee, J. H., and Jaffrey, S. R. (2019) m6A enhances the phase separation potential of mRNA. Nature 571, 424–428

55. Ditlev, J. A., Case, L. B., and Rosen, M. K. (2018) Who’s In and Who’s Out—Compositional Control of Biomolecular Condensates. Journal of Molecular Biology 430, 4666–4684

56. Sato, O., Sakai, T., Choo, Y. Y., Ikebe, R., Watanabe, T. M., and Ikebe, M. (2022) Mitochondria-associated myosin 19 processively transports mitochondria on actin tracks in living cells. Journal of Biological Chemistry 298, 101883

57. Shneyer, B. I., Ušaj, M., and Henn, A. (2016) Myo19 is an outer mitochondrial membrane motor and effector of starvation-induced filopodia. Journal of cell science 129, 543–556

58. Sun, H., Zhang, J., Ye, Q., Jiang, T., Liu, X., Zhang, X., Zeng, F., Li, J., Zheng, Y., Han, X., Su, C., and Shi, Y. (2023) LPGAT1 controls MEGDEL syndrome by coupling phosphatidylglycerol remodeling with mitochondrial transport. Cell Reports 42, 113214

59. Kasashima, K., Ohta, E., Kagawa, Y., and Endo, H. (2006) Mitochondrial Functions and Estrogen Receptor-dependent Nuclear Translocation of Pleiotropic Human Prohibitin 2*. Journal of Biological Chemistry 281, 36401–36410

