## Supplemental Figures for "Capturing the Hierarchically Assorted Protein-protein Interaction Modules of Mammalian Cell"

### Supplemental Figure S1

A

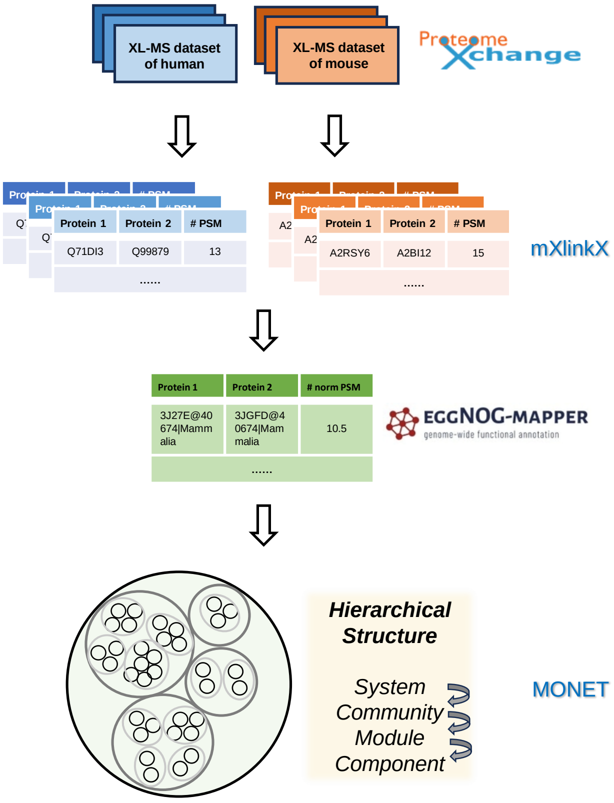


Step 1: Selecting the 7 MS-cleavable XL-MS datasets of Homo sapiens and Mus musculus from ProteomeXchange

Step 2: Searching the XL-MS datasets against target and decoy proteome database using mXlinkX and collecting the PSMs with FDR ≤ 0.01.

Step 3: Converting all repeatable crosslinked into mammalian ortholog interactions using EGGNOG-MAPPER. Normalizing the ortholog interaction weights among 7 datasets.

Step 4: Modulomic analysis of ortholog interactions to reveal their hierarchical organizations in cell.

**Supplemental Figure S1. Modulomic analysis pipeline.**

(A) The diagram represents the modulomic analysis pipeline of mammalian crosslinking mass spectrometry (XL-MS) datasets. Seven XL-MS datasets *in cell/tissue* and *in organello* level of Homo sapiens and Mus musculus were collected from ProteomeXchange, followed by crosslinked peptide searching through modified XlinkX (mXlinkX) using target-decoy strategy. The repeatable crosslinked peptides with FDR ≤ 0.01 were combined into mammalian using EGGNOG-MAPPER. Next, the interaction abundance of seven datasets were normalized using median normalization method. Finally, the hierarchal assorted structure of the interaction network was revealed by MONET-based modulomic analysis.

#### Supplemental Figure S2

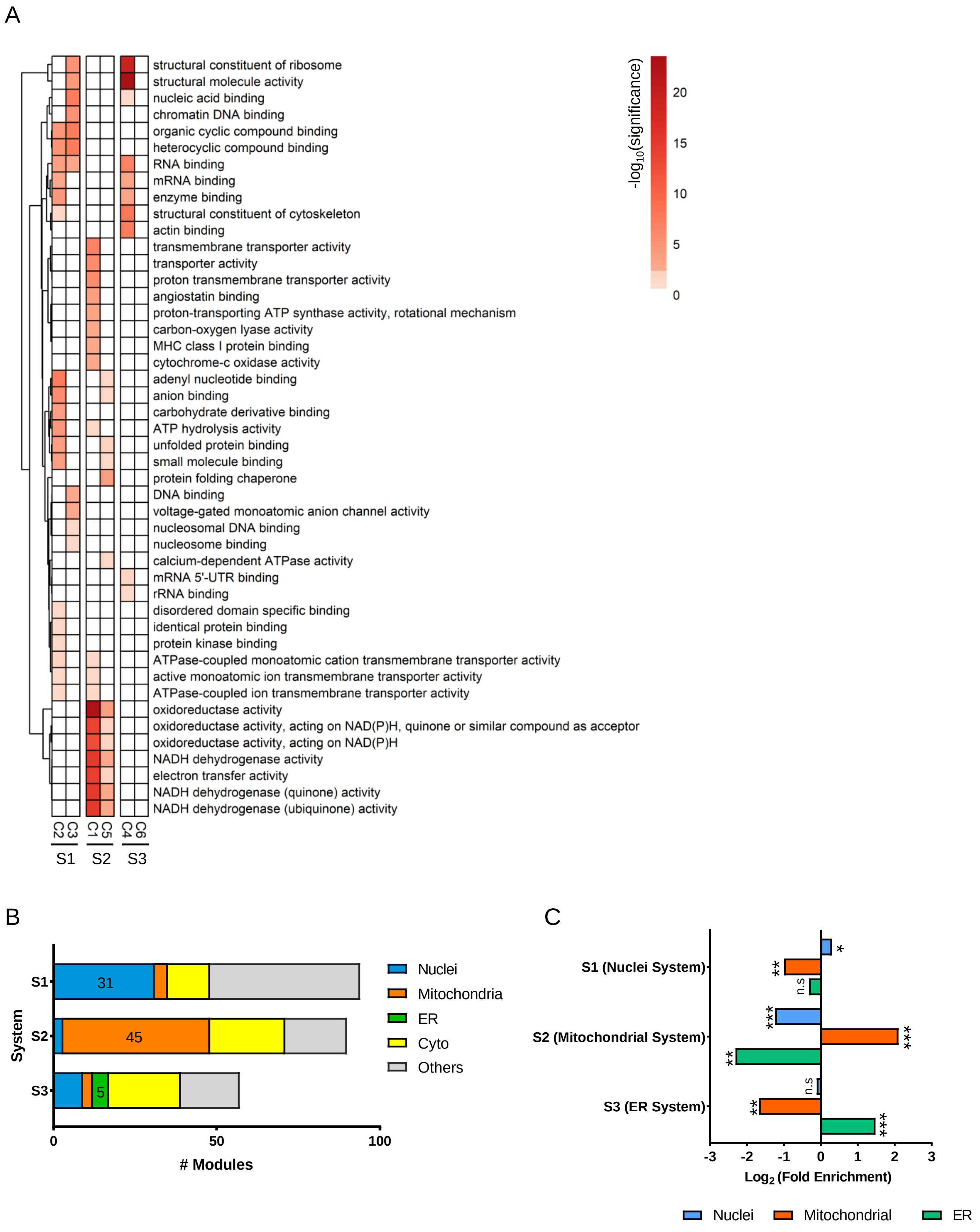

**Supplemental Figure S2. Characterization of communities and systems.**

(A) The heatmap of molecular function enrichment analysis of the ortholog proteins obtained each community. The communities are organized by systems. Row represents the name of GO term while column marks the different system and community of ortholog proteins. The color represents the negative number of base 10 logarithm of of each GO term. S1 - S3 stands for system 1 – 3. C1 – C6 stands for community 1 - 6. The data is listed in Table S3a-c.

(B) Bar Chat denotes distribution of ChatGPT-annotated subcellular localization of modules in three systems. The blue, orange, green, yellow and grey bar stands for nuclei, mitochondria, ER, cytosol and others, respectively. The data is listed in Table S2.

(C) Bar chart denotes the enrichment analysis results of subcellular localization of the highly interactive proteins in the three systems. The x axis represents the log2 ratio of fold enrichment. The blue, orange and green color stands for nuclei, mitochondrial and ER, respectively. Fisher exact test was applied. The n.s, *, ** and *** stands for $p > 0.05$, $p < 0.05$, $p < 0.01$ and $p < 0.001$, respectively. The data is listed in Table S2a.

#### Supplemental Figure S3

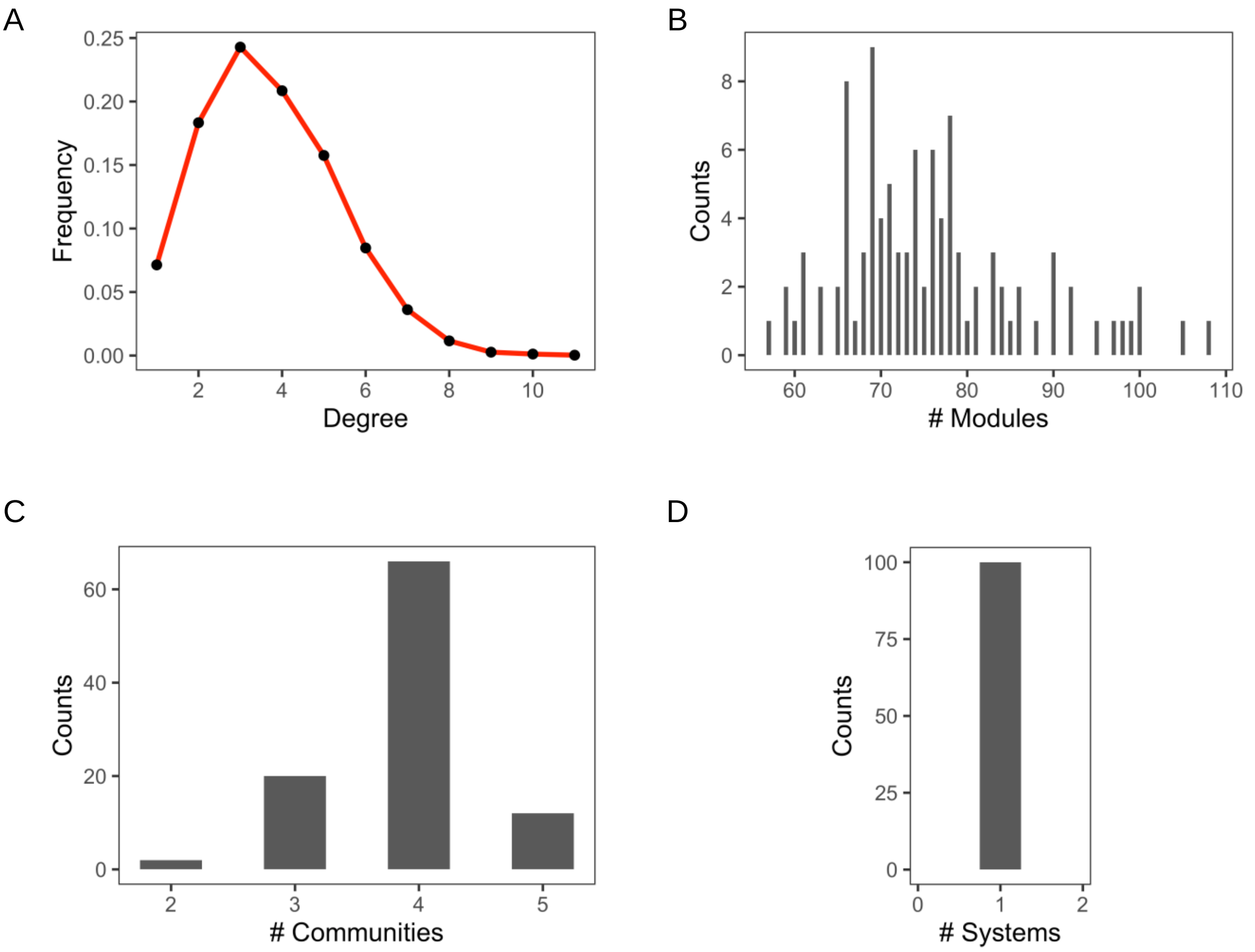

**Supplemental Figure S3. Modulome derived from randomly generated PPIs by Erdös-Rényi random network.**

(A) Line chart shows the protein degree distribution based on random PPI network. The frequency stands for the proportion of proteins with a specific number of degree. The random PPI network is an Erdös-Rényi random network generated by equivalent number of nodes (3377) and edges (4526) with the PPI network in this study (Table S1b; Erdös and Rényi, 1959).

(B - D) Histograms show the distribution of number of modules (B), communities (C) and systems (D) generated by 100 random tests. The y axis stands for the counts of tests with a specific number of modules (B), communities (C) and systems (D). The n indicates the 100 random tests. The details of random tests are described in Material and Methods.

### Supplemental Figure S4

A

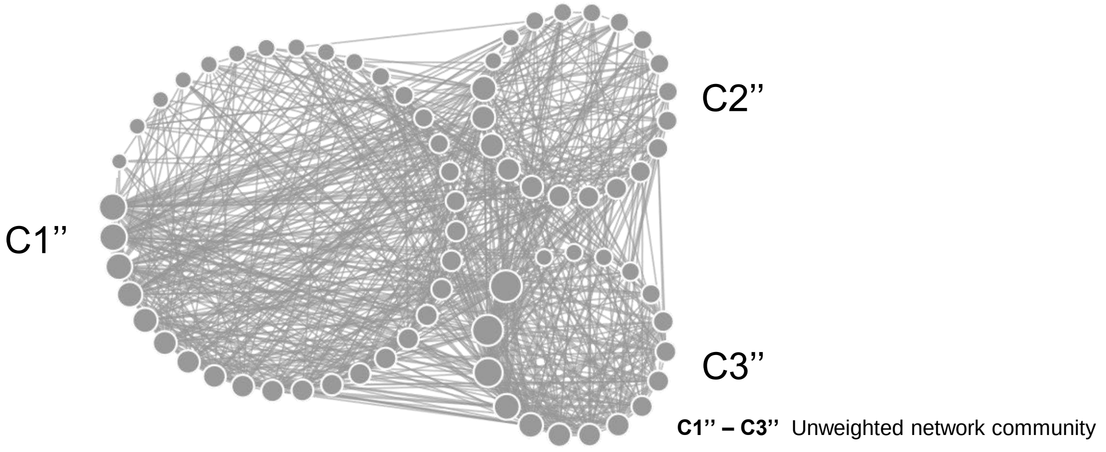


B

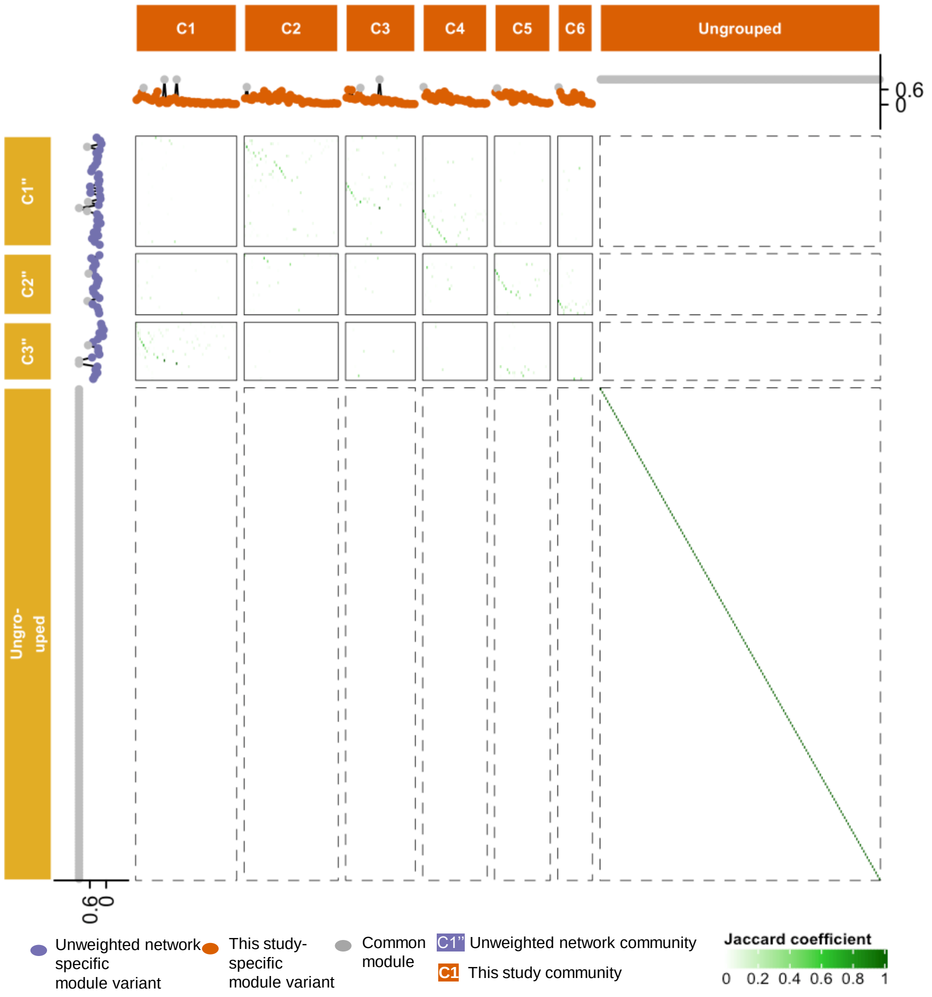


C

| | C1 | C2 | C3 | C4 | C5 | C6 |
|---|---|---|---|---|---|---|
| C1" | 0.04 | 0.35 | 0.25 | 0.22 | 0.01 | 0 |
| C2" | 0.05 | 0.1 | 0.02 | 0.08 | 0.26 | 0 |
| C3" | 0.53 | 0.01 | 0.02 | 0.02 | 0.02 | 0 |

**C1’’** Unweighted network community

**C1** This study community

**Jaccard coefficient**

0 0.2 0.4 0.6 0.8 1

**Supplemental Figure S4. Heatmap of modules and communities generated by unweighted PPI network (weight = 1) and this study.**

(A) The schematic topological graph represents communities generated by unweighted PPI network (3377 nodes, 4526 edges, weight = 1). The node and edge represents the modules and the interaction among modules (MMI, module-module interaction) within the cell graph, respectively. The higher order module of modules is defined as community. The size of the node represents the degree of the module. The thickness of the line represents the abundance of MMI, which is the count of PPIs between these two modules. The data are in Table S4a.

(B) The heatmap represents the comparison of modules generated by unweighted PPI network (3377 nodes, 4526 edges, weight = 1; rows; violet letters) and this study (columns; orange letters). The similarity between two modules is evaluated by Jaccard Coefficient (JC), which is determined by the number of identical components divided by the total number of unique components. The green color of the heatmap stands for the level of Jaccard Coefficient. The violet and orange dots stand for the unweighted network-specific module variants (JC <0.6), this study-specific module variants (JC<0.6), and common modules (JC>0.6), respectively (Table S4c). C1-C6 represents the community 1-6. Ungrouped are those modules that fail to be integrated into a community.

(C) The heatmap represents the comparison of communities generated by unweighted PPI network (rows; violet letters) and this study (columns; orange letters). The similarity between two modules is evaluated by Jaccard Coefficient (JC), which is determined by the number of identical components divided by the total number of unique components. The green color of the heatmap stands for the level of Jaccard Coefficient. C1-C6 represents the community 1-6. The data are in Table S4c.

### Supplemental Figure S5

A

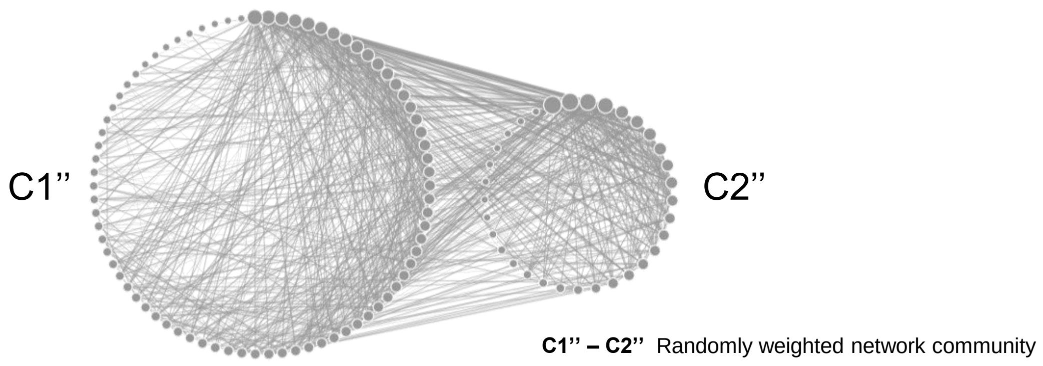


B

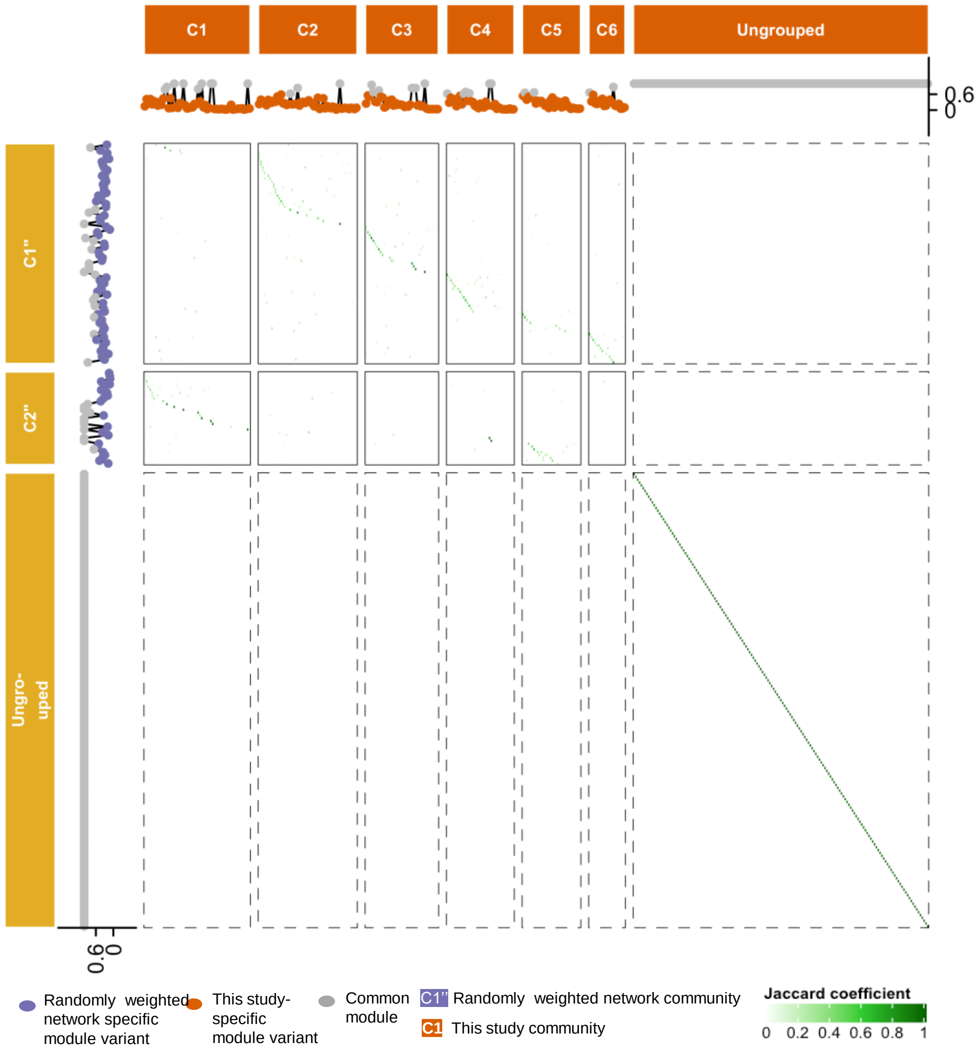


C

| | C1 | C2 | C3 | C4 | C5 | C6 |
|---|---|---|---|---|---|---|
| C1" | 0.06 | 0.29 | 0.2 | 0.2 | 0.07 | 0.11 |
| C2" | 0.44 | 0.04 | 0.01 | 0.02 | 0.18 | 0.01 |

C1'' Randomly weighted network community

C1 This study community

Jaccard coefficient

0 0.2 0.4 0.6 0.8 1

**Supplemental Figure S5. Heatmap of modules and communities generated by randomly weighted PPI network and this study.**

(A) The schematic topological graph represents communities generated by randomly weighted PPI network (3377 nodes, 4526 edges, randomly assign of the original PPI weight). The node and edge represents the modules (or module variants) and the interaction among modules (MMI, module-module interaction) within the cell graph, respectively. The higher order module of modules is defined as community. The size of the node represents the degree of the module. The thickness of the line represents the abundance of MMI, which is the count of PPIs between these two modules. The data are in Table S4b.

(B) The heatmap represents the comparison of modules generated by randomly weighted PPI network (3377 nodes, 4526 edges, randomly assign of the original PPI weight; rows; violet letters) and this study (columns; orange letters). The similarity between two modules is evaluated by Jaccard Coefficient (JC), which is determined by the number of identical components divided by the total number of unique components. The green color of the heatmap stands for the level of Jaccard Coefficient. The violet and orange dots stand for the random weighted network-specific module variants (JC <0.6), this study-specific module variants (JC<0.6), and common modules (JC>0.6), respectively (Table S4d). C1-C6 represents the community 1-6. Ungrouped are those modules that fail to be integrated into a community.

(C) The heatmap represents the comparison of communities generated by randomly weighted PPI network (rows; violet letters) and this study (columns; orange letters). The similarity between two modules is evaluated by Jaccard Coefficient (JC), which is determined by the number of identical components divided by the total number of unique components. The green color of the heatmap stands for the level of Jaccard Coefficient. C1-C6 represents the community 1-6. The data are in Table S4d.

### Supplemental Figure S6

**A**

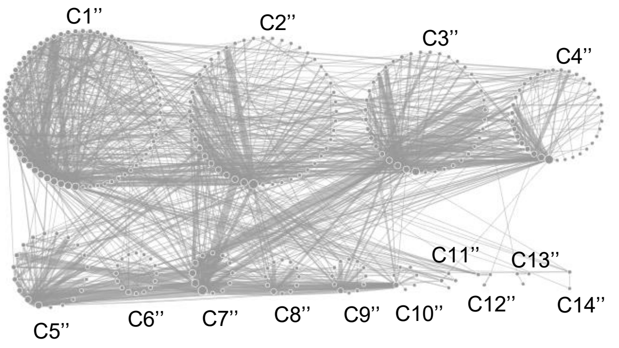


**C1” - C14”** Database community

**B**

| S1” | S2” |
|---|---|
| C1”, C2”, C3”, C4”, C6”, C8”, C9”, C11”, C12”, C13”, C14” | C5”, C7”, C10” |

S1” – S2” Database system
C1” – C14” Database community

**C**

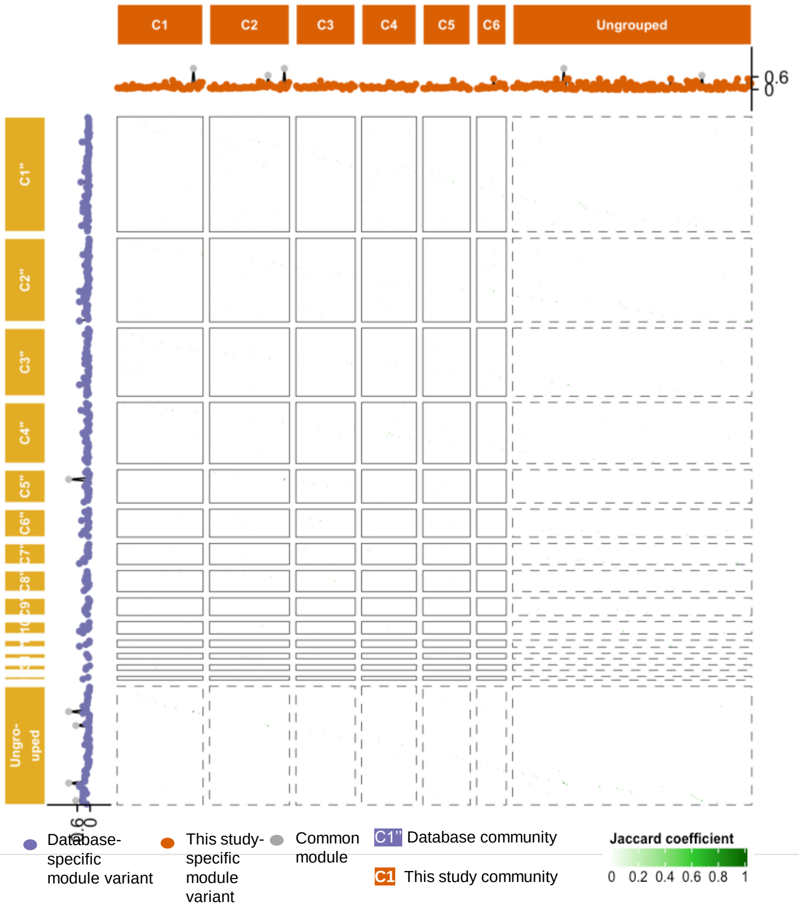


**D**

| | C1 | C2 | C3 | C4 | C5 | C6 |
|---|---|---|---|---|---|---|
| C1" | 0.05 | 0.09 | 0.07 | 0.1 | 0.07 | 0.07 |
| C2" | 0.04 | 0.07 | 0.09 | 0.06 | 0.04 | 0.04 |
| C3" | 0.17 | 0.04 | 0.02 | 0.02 | 0.06 | 0.02 |
| C4" | 0.04 | 0.06 | 0.02 | 0.05 | 0.05 | 0.03 |
| C5" | 0.02 | 0.05 | 0.07 | 0.03 | 0.02 | 0.02 |
| C6" | 0.01 | 0.02 | 0.05 | 0.01 | 0.02 | 0 |
| C7" | 0.05 | 0.03 | 0.05 | 0.1 | 0.04 | 0.03 |
| C8" | 0.01 | 0.02 | 0.03 | 0.02 | 0.01 | 0.01 |
| C9" | 0.01 | 0.01 | 0 | 0 | 0.01 | 0.01 |
| C10" | 0.01 | 0.01 | 0.01 | 0.01 | 0.01 | 0.02 |
| C11" | 0 | 0.01 | 0 | 0 | 0 | 0 |
| C12" | 0.01 | 0 | 0 | 0 | 0.02 | 0.01 |
| C13" | 0.01 | 0 | 0 | 0 | 0 | 0 |
| C14" | 0 | 0.01 | 0 | 0 | 0 | 0 |

**C1”** Database community
**C1** This study community

Jaccard coefficient
0 0.2 0.4 0.6 0.8 1

**Supplemental Figure S6. Comparison of modulome constructed by interactome derived from the databases and this study.**

(A) The schematic topological graph represents communities generated by database network (2316 nodes, 39196 edges). The node and edge represents the modules (or module variants) and the interaction among modules (MMI, module-module interaction) within the cell graph, respectively. The higher order module of modules is defined as community. The size of the node represents the degree of the module. The thickness of the line represents the abundance of MMI, which is the count of PPIs between these two modules. The data are in Table S4e.

(B) Table relationship between systems and communities conducted by database interactome. S1'' – S2'' stands for system 1 – 2. C1'' – C14'' stands for community 1 – 14 (Table S4f).

(C) The heatmap represents the comparison of modules generated by database network (2316 nodes, 39196 edges; rows; violet letters) and this study (columns; orange letters). The similarity between two modules is evaluated by Jaccard Coefficient (JC), which is determined by the number of identical components divided by the total number of unique components. The green color of the heatmap stands for the level of Jaccard Coefficient. The violet and orange dots stand for the random weighted network-specific module variants (JC <0.6), this study-specific module variants (JC<0.6), and common modules (JC>0.6), respectively (Table S4f). C1-C6 represents the community 1-6. Ungrouped are those modules that fail to be integrated into a community.

(D) The heatmap represents the community comparison of communities between database (rows; violet letters) and this study (columns; orange letters). The similarity between two modules is evaluated by Jaccard Coefficient (JC), which is determined by the number of identical components divided by the total number of unique components. The green color of the heatmap stands for the level of Jaccard Coefficient. C1-C6 represents the community 1-6. The data are in Table S4f.

### Supplemental Figure S7

**A**

| | C2 | C3 | C1 | C5 | C4 | C6 |
|---|---|---|---|---|---|---|
| C1" | 0.19 | 0.07 | 0.04 | 0.09 | 0.21 | 0.08 |
| C2" | 0.07 | 0.21 | 0.08 | 0.05 | 0.05 | 0 |
| C3" | 0.11 | 0.13 | 0.74 | 0.45 | 0.05 | 0.02 |
| C4" | 0.23 | 0.09 | 0.29 | 0.3 | 0.06 | 0.01 |
| C6" | 0.11 | 0.09 | 0.04 | 0.04 | 0.07 | 0.12 |
| C8" | 0.16 | 0.24 | 0.1 | 0.07 | 0.09 | 0 |
| C9" | 0.04 | 0 | 0 | 0.03 | 0 | 0 |
| C11" | 0 | 0 | 0 | 0 | 0 | 0 |
| C12" | 0 | 0 | 0 | 0 | 0 | 0 |
| C13" | 0 | 0 | 0.02 | 0.02 | 0 | 0 |
| C14" | 0 | 0 | 0 | 0 | 0 | 0 |
| C5" | 0.11 | 0.18 | 0.1 | 0.07 | 0.05 | 0 |
| C7" | 0.13 | 0.28 | 0.21 | 0.2 | 0.22 | 0 |
| C10" | 0.03 | 0 | 0.01 | 0.02 | 0.03 | 0 |

**B**

| | C2 | C3 | C1 | C5 | C4 | C6 |
|---|---|---|---|---|---|---|
| C1" | 0.05 | 0.02 | 0.01 | 0.03 | 0.1 | 0.01 |
| C2" | 0.02 | 0.07 | 0.05 | 0.02 | 0.08 | 0.01 |
| C3" | 0.1 | 0.05 | 0.51 | 0.23 | 0.06 | 0 |
| C4" | 0.12 | 0.04 | 0.04 | 0.12 | 0.13 | 0 |
| C6" | 0.09 | 0.07 | 0.03 | 0.05 | 0.12 | 0 |
| C8" | 0.03 | 0.04 | 0.02 | 0.01 | 0.02 | 0.07 |
| C9" | 0.03 | 0 | 0 | 0.02 | 0.02 | 0 |
| C11" | 0 | 0 | 0 | 0 | 0 | 0 |
| C12" | 0.02 | 0.02 | 0.04 | 0.02 | 0.01 | 0 |
| C13" | 0.02 | 0 | 0 | 0.03 | 0 | 0 |
| C14" | 0 | 0 | 0 | 0 | 0.01 | 0 |
| C5" | 0.1 | 0.12 | 0.06 | 0.02 | 0.13 | 0 |
| C7" | 0.06 | 0.27 | 0.08 | 0.04 | 0.45 | 0 |
| C10" | 0 | 0.06 | 0.07 | 0.02 | 0.11 | 0 |

**C**

| | C2 | C3 | C1 | C5 | C4 | C6 |
|---|---|---|---|---|---|---|
| C1" | 0.25 | 0.02 | 0.01 | 0.05 | 0.1 | 0 |
| C2" | 0.12 | 0.15 | 0 | 0.02 | 0.06 | 0 |
| C3" | 0.03 | 0 | 0.48 | 0.17 | 0 | 0 |
| C4" | 0.14 | 0.03 | 0 | 0.11 | 0.08 | 0 |
| C6" | 0.06 | 0.03 | 0 | 0.06 | 0.03 | 0 |
| C8" | 0 | 0 | 0 | 0 | 0 | 0 |
| C9" | 0 | 0 | 0 | 0 | 0 | 0 |
| C11" | 0 | 0 | 0 | 0 | 0 | 0 |
| C12" | 0 | 0 | 0.08 | 0.04 | 0 | 0 |
| C13" | 0 | 0 | 0 | 0 | 0 | 0 |
| C14" | 0 | 0 | 0 | 0 | 0 | 0 |
| C5" | 0.09 | 0.1 | 0 | 0.02 | 0.1 | 0 |
| C7" | 0.09 | 0.27 | 0 | 0.03 | 0.3 | 0 |
| C10" | 0.02 | 0.04 | 0.03 | 0 | 0.04 | 0 |

**Supplemental Figure S7. Comparison of the biological role of modulome constructed by interactome derived from the databases and this study.**

(A – C) The heatmap shows the comparison of cellular component (A), biological process (B), and molecular function (C) analysis of proteins derived from database (rows; violet letters) and this study (columns; orange letters) communities. The top 20% of most interactive proteins (highest protein degree) in each community were used for the analysis. The similarity among the communities is evaluated by the Jaccard Coefficient (JC) of related GO terms. The green color of the heatmap stands for the level of the Jaccard Coefficient (Table S4f).

### Supplemental Figure S8

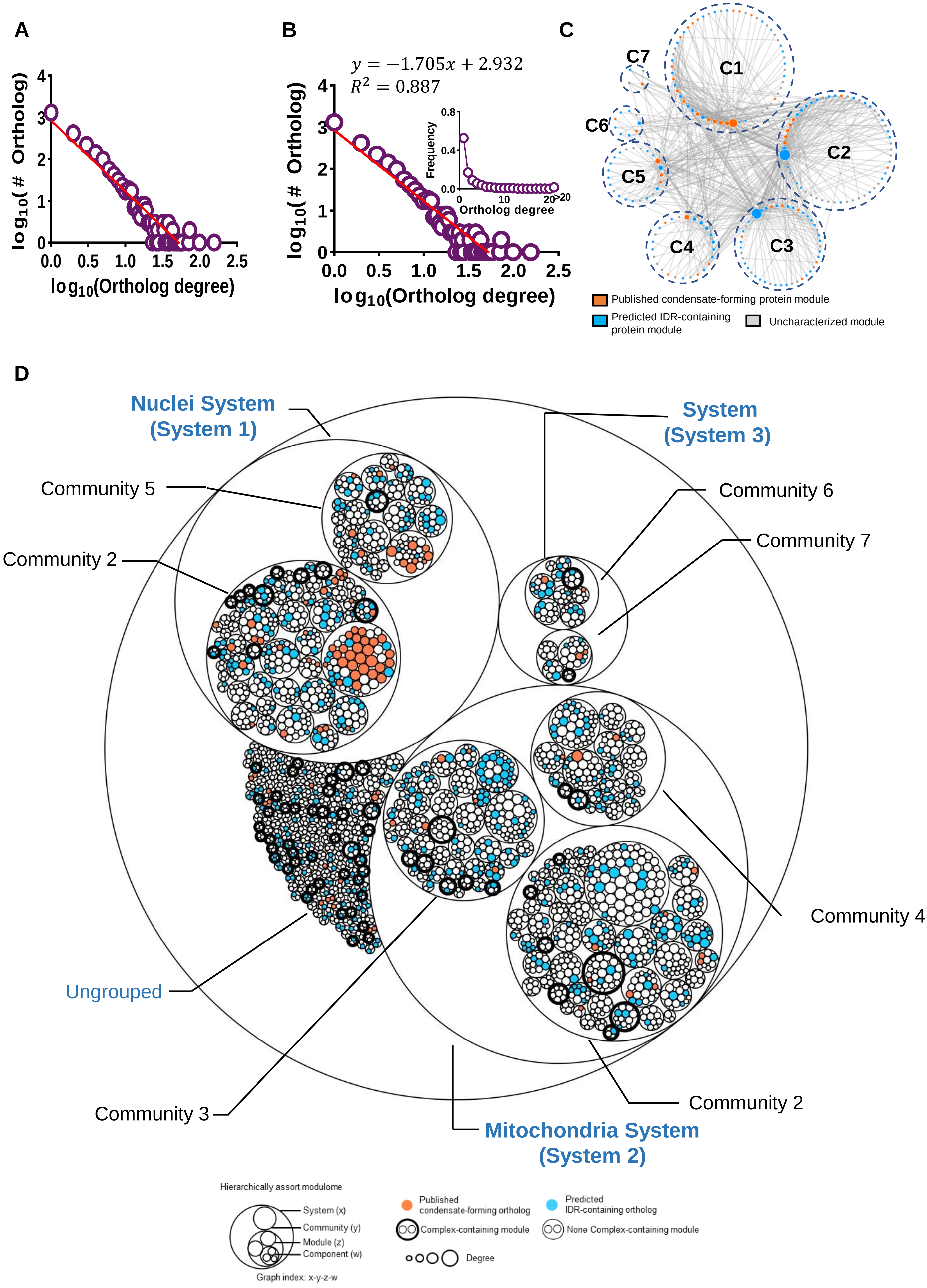

**Supplemental Figure S8. Modulome constructed by XL-MS datasets derived from the original papers.**

(A) Pie chart represents the distribution of hetero- (purple; 3969 interactions) and homo-ortholog interaction (grey; 3239 interactions) combined from repeatable crosslinks with FDR ≤ 0.01 (Table S4g).

(B) Line diagram (left-bottom panel) shows the distribution of degree 2476 ortholog proteins derived 3239 hetero-ortholog interactions (Table S4g). The power law equation of the fitting curve (right-top panel) describes the correlation in between the total number of ortholog protein (2476) and the specific number of degree of an ortholog protein. $R^2$ represents the coefficient of the curve determination (Table S4g).

(C) Network diagram represents seven communities present in mammalian cell. The node and edge represents the module and the interaction among modules, which is defined as module-module interaction (MMI), respectively. The higher order of modules (module of module) is defined as community. The size of the node represents the degree of module. The orange, light blue and light grey node represents the published condensate-forming, the predicted IDR-containing and uncharacterized module, respectively. The thickness of line represents the abundance of MMI. C1 – C7 stands for community 1 - 7. The data is listed in (Table S4h).

(D) Circle plot denotes the hierarchically assorted modulome containing system, community, modules and protein components, generated by multiple rounds of modulomic analysis of ortholog interaction network. Circle denotes the ortholog proteins. The size of circle denotes the protein degree. The orange color, blue color, and white color filled circle represents published condensate-forming, IDR-containing and uncharacterized ortholog protein, respectively. The bold outline of circle indicates a module which is overlapped with well-reported protein complex. The outline of grouped circles (ortholog protein components) denotes a module. The outline of grouped modules denotes a community. The outline of grouped community denotes a system. Ungrouped are those modules that fail to be integrated into a community. The biological character of each system is determined by GO enrichment analysis. The data is listed in Table S4h.

#### Supplemental Figure S9

**A**

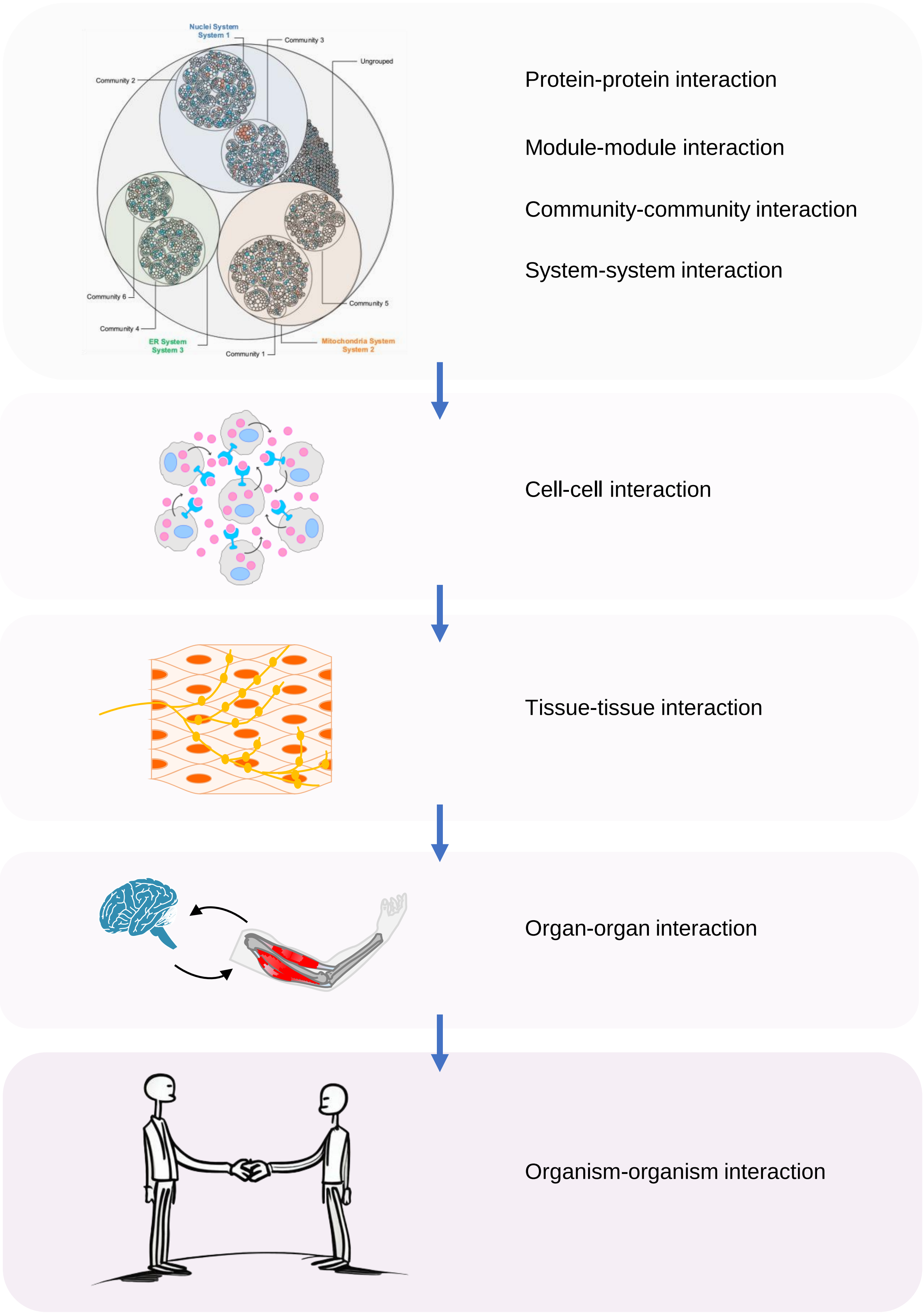

**Supplemental Figure S9. Modulome of organism.**

(A) Diagram represent hierarchically assorted modulome of organism.
